# Regeneration in calcareous sponge relies on ‘purse-string’ mechanism and the rearrangements of actin cytoskeleton

**DOI:** 10.1101/2022.11.02.514829

**Authors:** Kseniia V. Skorentseva, Fyodor V. Bolshakov, Alina A. Saidova, Andrey I. Lavrov

**Affiliations:** Koltzov Institute of Developmental Biology of Russian Academy of Sciences, Moscow, Russia; Pertsov White Sea Biological Station, Biological Faculty, Lomonosov Moscow State University, Moscow, Russia; Department of Cell Biology and Histology, Biological Faculty, Lomonosov Moscow State University, Moscow, Russia; Engelhardt Institute of Molecular Biology, Russian Academy of Sciences, Moscow, Russia

**Keywords:** actin filaments, actomyosin cable, epithelisation, morphogenesis, Porifera, Calcarea

## Abstract

The crucial step in any regeneration process is epithelization, i.e. the restoration of epithelium structural and functional integrity. Epithelialization requires cytoskeletal rearrangements, primarily of actin filaments and microtubules. Sponges (phylum Porifera) are early branching metazoans with pronounced regenerative abilities. Calcareous sponges have a unique step during regeneration: formation of a temporary structure, regenerative membrane which initially covers a wound. It forms due to the morphallactic rearrangements of exo- and choanoderm epithelial-like layers. The current study quantitatively evaluates morphological changes and characterises underlying actin cytoskeleton rearrangements during regenerative membrane formation in asconoid calcareous sponge *Leucosolenia variabilis*, through a combination of time-lapse imaging, immunocytochemistry, and confocal laser scanning microscopy. Regenerative membrane formation has non-linear stochastic dynamics with numerous fluctuations. The pinacocytes at the leading edge of regenerative membrane form a contractile actomyosin cable. Regenerative membrane formation either depend on its contraction or being coordinated through it. The cell morphology changes significantly during regenerative membrane formation. Exopinacocytes flatten, their area increases, while circularity decreases. Choanocytes transdifferentiate into endopinacocytes, losing microvilli collar and flagellum. Their area increases and circularity decreases. Subsequent redifferentiation of endopinacocytes into choanocytes is accompanied by inverse changes in cell morphology. All transformations are based on actin filament rearrangements similar to those characteristic of higher metazoans. Altogether, we provide here a qualitative and quantitative description of cell transformations during reparative epithelial morphogenesis in a calcareous sponge.

**Summary statement:** First detailed description of actin cytoskeleton rearrangements during wound healing in calcareous sponge. ‘Purse-string’ mechanism presumably involved. Cytoskeletal rearrangements resemble those characteristic of Eumetazoans.

## Introduction

Cell motility plays a key role in embryogenesis, wound healing, and immune response. Changes in cell morphology and migration potential occur due to cytoskeletal rearrangements, in particular, due to coordinated functioning of an actomyosin complex and the presence of cell adhesions, anchoring the cell to the substrate or binding it to the other cells (Babbin *et al*., 2009; Blanchoin *et al*., 2015).

Healing epithelial wounds is of crucial importance to restoring the internal environment of any animal. It was considered for a long time that embryonic wounds are healed by a special, actin “purse-string” mechanism (Martin and Lewis, 1992; Brock *et al*., 1996): cells on the leading edge of the wound are forming a continuous actomyosin ring and its contraction pulls wound edges together. On the contrary, epithelial wounds in adult organisms were considered to be healed by extension of lamellipodia from cells at the leading edge followed by their collective migration into the wound (Pang *et al*., 1978; Zahm *et al*., 1991; Kretschmer *et al*., 2017). During such a process, epithelial cells lose their apical/basolateral polarity and flatten (Bement *et al*., 1993). However, the new data brought the problem to a new level of complexity. To date, two major mechanisms, purse-string and collective cell migration, are considered to interact with each other through the crosstalk of signalling pathways (Anon *et al*., 2012; Brugués *et al*., 2014; Kuipers *et al*., 2014), in particular through the functioning of the Rho family of GTPases (Nobes and Hall, 1995; Fenteany *et al*., 2000; Tamada *et al*., 2007).

Sponges (phylum Porifera) is one of the most basal metazoan group, which had long independent evolution (Simion *et al*., 2017). Sponges have a distinctive body plan and possess unique regenerative capacities ranging from wound healing to complete restoration of functional organism from dissociated cells (Lavrov and Kosevich, 2014; Ereskovsky *et al*., 2021). The mechanisms of regeneration vary significantly among sponge clades. The regeneration in Demospongiae relies on various transformations of individual cells: dedifferentiation, transdifferentiation, epithelial-mesenchymal transition (EMT) and mesenchymal-epithelial transition (MET) (Alexander *et al*., 2015; Borisenko *et al*., 2015; Ereskovsky *et al*., 2020). In contrast, regeneration in Calcarea is based on coordinated interaction of the epithelial cell layers with no contribution of proliferation to wound healing process (Lavrov *et al*., 2018; Caglar *et al*., 2021). This process requires cell transdifferentiation and epithelial transformations followed by loss of characteristic cell features (Lavrov *et al*., 2018; Ereskovsky and Lavrov, 2021).

Essential studies on calcareous sponge regeneration were carried out mainly using light and electron microscopy (Korotkova, 1961, 1963; Padua and Klautau, 2016; Ereskovsky et al., 2017; Lavrov *et al*., 2018). At the site of the injury, cells of the body wall form a two-layered semi-transparent membrane, regenerative membrane (RM). The process has two phases: the membrane formation itself and its transformation into the normal body wall, i.e., choanoderm formation on its inner side, porocytes development, and spiculogenesis. Regeneration proceeds due to the rearrangements and migration of the epithelial-like cell layers of the pinaco- and choanoderm without loss of cell-cell adhesions or proliferation (Korotkova, 1961, 1963; Lavrov *et al*., 2018; Caglar *et al*., 2021).

Currently, data on the structure of actin filaments in sponge cells are fragmentary and obtained primarily for freshwater sponges (Pavans de Ceccatty, 1981, 1986; Wachtmann *et al*., 1990; Gaino and Magnino, 1999), and the most detailed studies describe the role of the actin cytoskeleton in the formation of cell adhesions (Miller *et al*., 2018; Mitchell and Nichols, 2019; Green *et al*., 2020). The role of cytoskeletal rearrangements in morphogenetic processes during embryogenesis and regeneration in sponges remains poorly elucidated.

In the current study, we examined the general dynamics and rearrangements of actin filaments in main cell types during body wall regeneration in asconoid calcareous sponge *Leucosolenia variabilis* using immunocytochemistry, confocal laser scanning microscopy, and live imaging time-lapse imaging. Our study reveals cytoskeleton rearrangements underlying transformations of epithelial-like cell layers (exopinacoderm and choanoderm) and formation of contractile actomyosin cable on the leading edge of the wound. We show significant changes in cell morphology during this epithelial morphogenesis. We also studied the morphology and dynamic parameters of the individual mesohyl cells in the RM, thus providing new data on epithelial morphogenesis as the main force of regeneration in calcareous sponges.

## Materials and methods

### Sampling and surgical operation

Specimens of *Leucosolenia variabilis* Haeckel, 1870 (Calcarea, Leucosolenida) were collected during summer seasons 2019-2022 in Kandalaksha Bay, the White Sea, in the vicinity of the Pertsov White Sea Biological Station of Lomonosov Moscow State University (66°34’N, 33°08’E). The specimens were collected in the upper subtidal zone (0 – 2 m) during low tides and stored in the flow-through aquarium for no longer than 10 days.

As a regeneration model in this study, we used ring-shaped body fragments from oscular tubes. The operation procedures were performed in 0.22-μm-filtered seawater (FSW). Sponges were rinsed from debris, oscular tubes were cut off and collected. Each oscular tube was transversely cut into ring-shaped body fragments 1-3 mm width (the osculum rims were excluded). All surgical operations were performed manually under a stereomicroscope using Castroviejo scissors. Body fragments were maintained in Petri dishes with 5 ml FSW at the physiological temperature 8-12°C (Fig. S1). Half of the FSW medium was changed every 12 hours. Regular observations were made to prevent contamination and assess the stage of regeneration.

### Time-lapse imaging

To visualise regenerative membrane growth and track individual mesohyl cells in it, time-lapse imaging was performed using a Nikon TI-S inverted microscope equipped with a digital camera ToupCam U3CMOS05100KPA (Touptek Photonics, China) (application ToupView v. 3.7) and cooling microscopic stage. For recording, body fragments were maintained in glass bottom Petri dishes with 5 ml FSW at temperatures of 10-12°C. The membrane growth was filmed with an objective 20x LWD Achromat 0.40 Ph1 ADL WD 3.1 ∞/1.2 OFN25 (Nikon Instruments Inc, USA) with an additional lens of 0.5x with 30 sec intervals between frames. For cell tracking, an objective 40x ELWD S Plan Fluor 0.60 Ph2 ADM WD 3.6-2.8 ∞/0-2 OFN22 (Nikon Instruments Inc.) with an additional lens 0.5x was used; interval between frames was 15 seconds (Lavrov and Ereskovsky, 2020).

To perform cell tracking, the MTrackJ plugin in ImageJ v.1.53c (National Institute of Health, Bethesda) was used. We calculated total track distance, total displacement (the length of line connecting first and last track point), the average cell velocity (dividing the total track distance divided by the acquisition time) and migration efficiency (dividing the total displacement by the track length). The measurements were performed at two stages of regeneration: during the active membrane growth (12-24 hours past operation, or hpo) and in full membrane during choanocyte redifferentiation (24-48 hpo).

The following programs were used to create time-lapse films: ImageJ v.1.53c, Adobe Photoshop Lightroom Classic V.10.3 (Adobe Inc., USA), VirtualDub (https://www.virtualdub.org), Adobe Premier Pro V.22.1.2 (Adobe Inc., USA) and Handbrake (https://github.com/HandBrake/HandBrake).

### Immunostaining

Four fixation methods were used: 4% paraformaldehyde (Carl Roth 0335.2) in phosphate-buffered saline (PBS, Eco-Servis B-60201); 4% paraformaldehyde (Carl Roth 0335.2) in FSW; 2.5% glutaraldehyde (EMS 16220) in PBS at 4°C for at least 4 hours; ice-cold methanol (Merck 106008) for 1 hour followed by 4% PFA in PBS at 4°C overnight. All fixation methods provided comparable results.

After fixation, the samples were rinsed with PBS three times. To prevent autofluorescence of glutaraldehyde, samples were incubated in three changes of 1-3% sodium borohydride (NaBH4, Panreac 163314) for an hour. The samples were then blocked with a solution of 1% bovine serum albumin (BSA, MP Biomedicals 0216006980), 0.1% cold-water fish skin gelatine (Sigma-Aldrich G7041), 0.5% Triton X-100 (Sigma-Aldrich T8787), 0.05% Tween-20 (Sigma-Aldrich P1379) in PBS and incubated overnight at 4°C in primary antibodies: anti-human actin-β primary monoclonal antibody produced in mouse (1:100; Bio-Rad Laboratories Inc. MCA57766A) and non-muscle myosin II produced in rabbit (1:100, Sigma-Aldrich M8064). In some samples, Phalloidin FITC (1:200, Sigma-Aldrich P5282) was used instead of primary antibodies to reveal actin filaments, since staining patterns appeared equal. After incubation with antibodies, the samples were rinsed three times in the blocking solution and incubated for 4 hours in goat polyclonal secondary antibody to mouse IgG conjugated with AlexaFluor 555 (1:100, Invitrogen A31570) and donkey polyclonal secondary antibody to rabbit IgG conjugated with AlexaFluor 647 (1:100, Invitrogen A10040). Finally, the samples were rinsed three times in PBS and stained with 2 μg ml^-1^ 4’,6-diamidino-2-phenylindole (DAPI, Acros 202710100) in PBS for 1 hour. Rinsed specimens were mounted in 90% glycerol (MP Biomedicals 193996) with 2.5% 1,4-23 diazabicyclo[2.2.2]octane (DABCO, Sigma-Aldrich D27802) and examined with a CLSM Nikon A1 (Nikon, Shinagawa, Japan) using lasers with 405, 546 and 647 nm wavelength.

### Image Processing and Statistical Analysis

The obtained Z stacks (step 200-300 nm) were processed with ImageJ v.1.53c to describe actin structures in different cell types and define the morphological parameters of the cells: area, circularity, and aspect ratio. For intact exopinacocytes, the cell body and cytoplasmic plates were analysed separately. Circularity was calculated with the following formula: 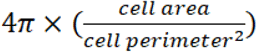 (1). A high circularity value (tending to 1.0) indicates fewer cell protrusions (such as filopodia or lamellipodia) and a low value (tending to zero) – the unevenness of the cell edges. The aspect ratio (AR) was calculated using the following formula: 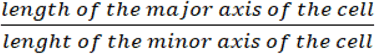 (2). It illustrates how elongated the cell is: the higher values correspond to narrower and longer cells. To obtain these parameters, cell images were outlined using the ‘Freehand selection’ tool and “ROI manager”. The raw data is provided in the Tables S1, S2.

The digital photos were processed in ImageJ v.1.53c. Gaussian blur, changing the brightness-contrast parameters and background subtraction were applied to 16-bit images for noise reduction and contrast enhancement. This processing sequence was used both for time-lapse imaging and confocal images.

To compare the morphological parameters of different cell types and the dynamic parameters of mesohyl cells, the following statistical tests were used: Shapiro-Wilk test, Kruskal-Wallis test by ranks, Mann-Whitney *U* test and Dunn’s multiple comparisons test with Benjamini-Hochberg correction. Firstly, the datasets were tested for normality and outliers were excluded using the ROUT method (Q = 5%) in Prism 8.0.1 (GraphPad Software, USA). Comparisons were handled using an appropriate statistical test (parametric/nonparametric, multiple comparisons/not multiple). A test used is indicated in each case individually. These statistical procedures and data visualisation were performed in RStudio version 4.0.3 (RStudio, USA). All the data are expressed as mean ± s.e.m. (standard error of mean). The significance level was 0.05.

## Results

### General observations on the formation of the regenerative membrane

*Leucosolenia variabilis* has an asconoid body plan. Its body wall has a thickness of 20–30 μm and contains three cell layers: the outer exopinacoderm, the inner choanoderm, and intermediate mesohyl, which contains a few migrating cells and the extracellular matrix (Fig. 1A, B).

**Figure 1.**
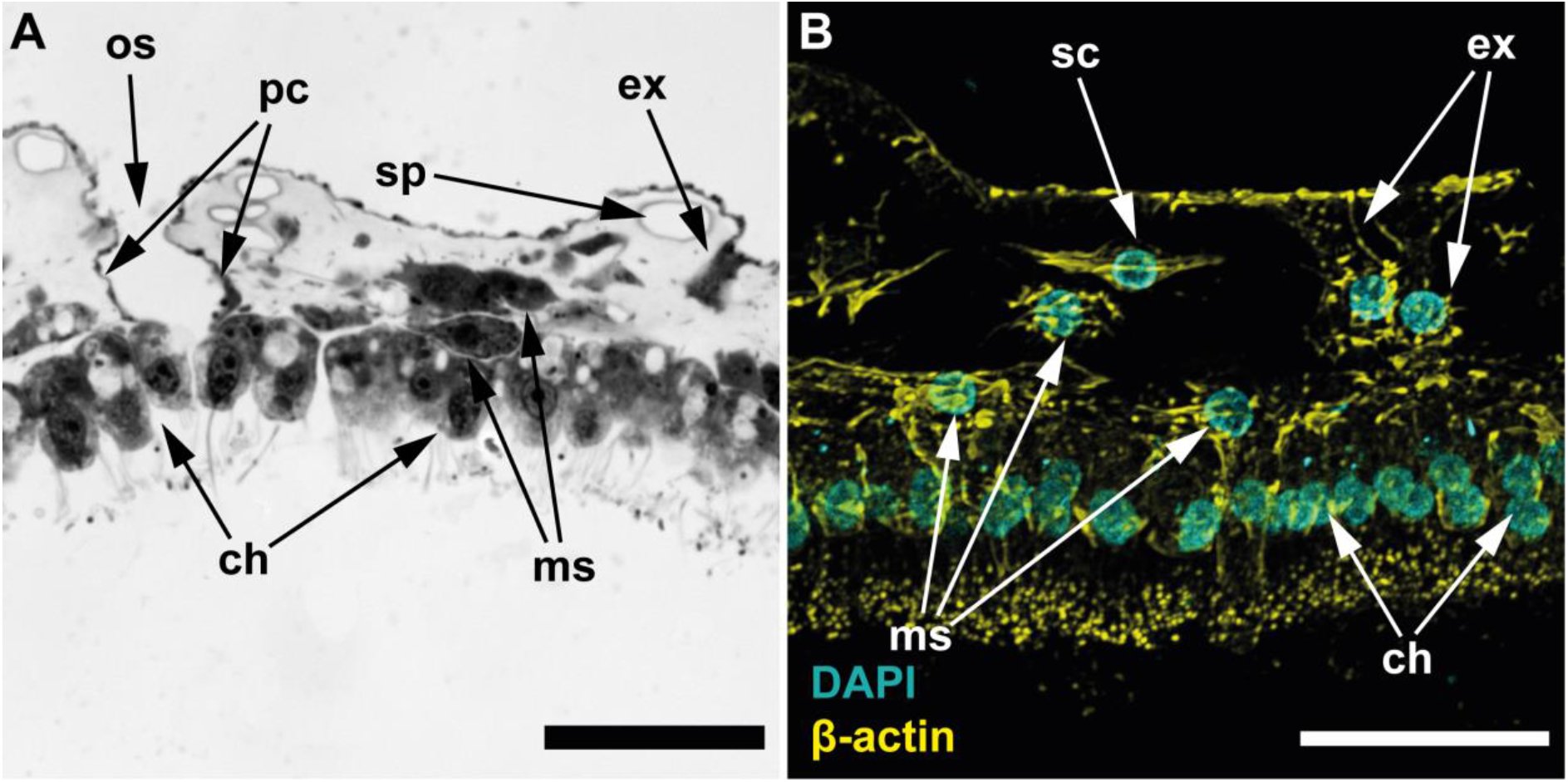
Histology of *Leucosolenia variabilis* body wall, cross-section through the middle part of the oscular tube. (A) semi-thin section, light microscopy; (B) CLSM, maximum intensity projection of several focal planes; cyan – DAPI, cell nuclei; yellow – cytoplasmic β-actin isoform, actin filaments. ch, choanoderm; ex, exopinacocyte; ms, mesohyl cell; os, ostium; pc, porocyte; sc, sclerocyte; sp, spicule. Scale bars: (A–B) – 20 μm.

Ring-shaped body fragments obtained after the operation have two wound areas located in the cut planes. The subsequent regeneration process leads to the formation of new body walls, which close the orifices and are perpendicular to the intact walls of the body fragment (Fig. S1).

The regeneration occurs through the formation of a temporary structure, the regenerative membrane (RM). RM appears at ~12-24 hpo on the edge of the wound and gradually expands towards its centre, until forming a continuous structure (Fig. 2A-E; Movie S1). During growth, RM has a thickness of 10-15 μm and consists of three layers: the external exopinacoderm, the internal endopinacoderm, and very narrow mesohyl in between (Movie S1). As shown previously, endopinacocytes of RM appear in the wound area through the transdifferentiation of the choanocytes (Lavrov *et al*., 2018). The exo- and endopinacocytes adhesions at the growing rim of RM isolate the mesohyl layer from the external environment.

**Figure 2.**
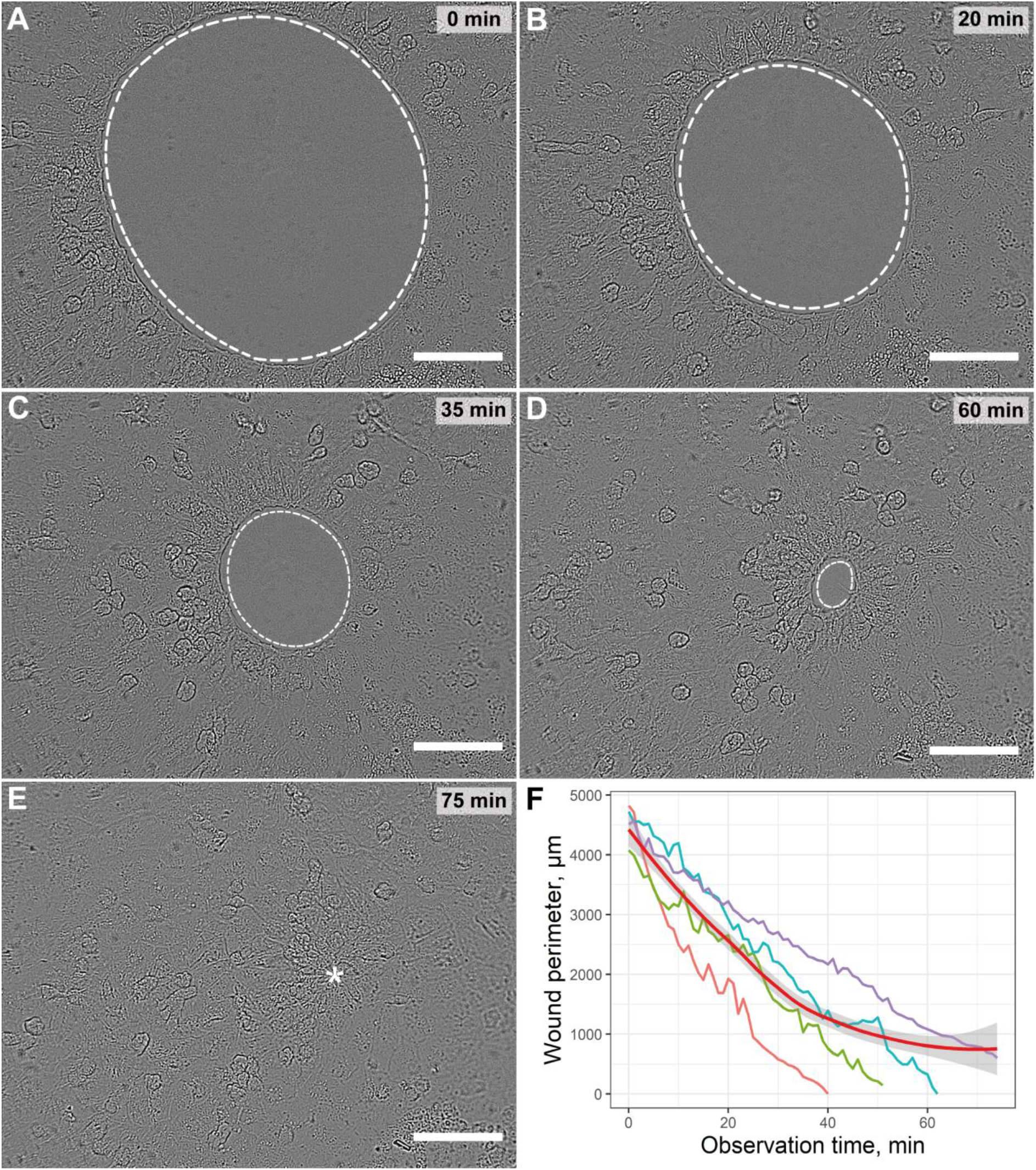
Dynamics of regenerative membrane (RM) formation prior to full closure (~24 hpo). (A-E) RM closure, light microscopy, frames from time-lapse imaging; (F) line plot of closure dynamics (n=4), red line is the trend line with a confidence interval (grey background). White dotted lines mark the leading edge of the RM; asterisk mark the RM closure point. Scale bars (A-E) – 50 μm.

The complete RM closing the wound orifice usually forms at ~48 hpo. However, already at ~24-30 hpo, the transformation of RM into the body wall begins. This process includes the redifferentiation of endopinacocytes to choanocytes at the internal side of RM (Lavrov *et al*.,2018), appearance of porocytes penetrating RM, and synthesis of new spicules in the RM mesohyl. All these processes begin in the peripheral part of RM and gradually proceed in the centripetal direction. The complete transformation of RM into the body wall ends at 120-144 hpo for body wall cuts (Lavrov *et al*., 2018), although tracking the process to its end in our model was beyond the scope of the study.

The formation and transformation of the RM rarely appear to be symmetrical: different sides of the RM grow and turn into the body wall at a varying rate. Moreover, time-lapse visualisations clearly show that RM is under significant mechanical tension since sometimes it rips on the leading edge (Movie S2). To describe RM growth dynamics, we analysed it 1-2 hours before a complete closure over a wound plane. The RM growth appears to be nonlinear process with fluctuations. The obtained data demonstrate also a varying regeneration speed rate between individual specimens (time spent on closing the hole with a perimeter of 4-5 mm varied from 40 to 80 minutes) (Fig. 2F).

### Mesohyl cell dynamics in the regenerative membrane

Time-lapse visualisation allows us to track individual mesohyl cells in the RM during its growth and transformation (Fig. 3; Movie S1, S2). They can be approximately divided into three categories based on their appearance and behaviour, but we are not able to unambiguously match the described types with cell subtypes in intact mesohyl tissue.

**Figure 3.**
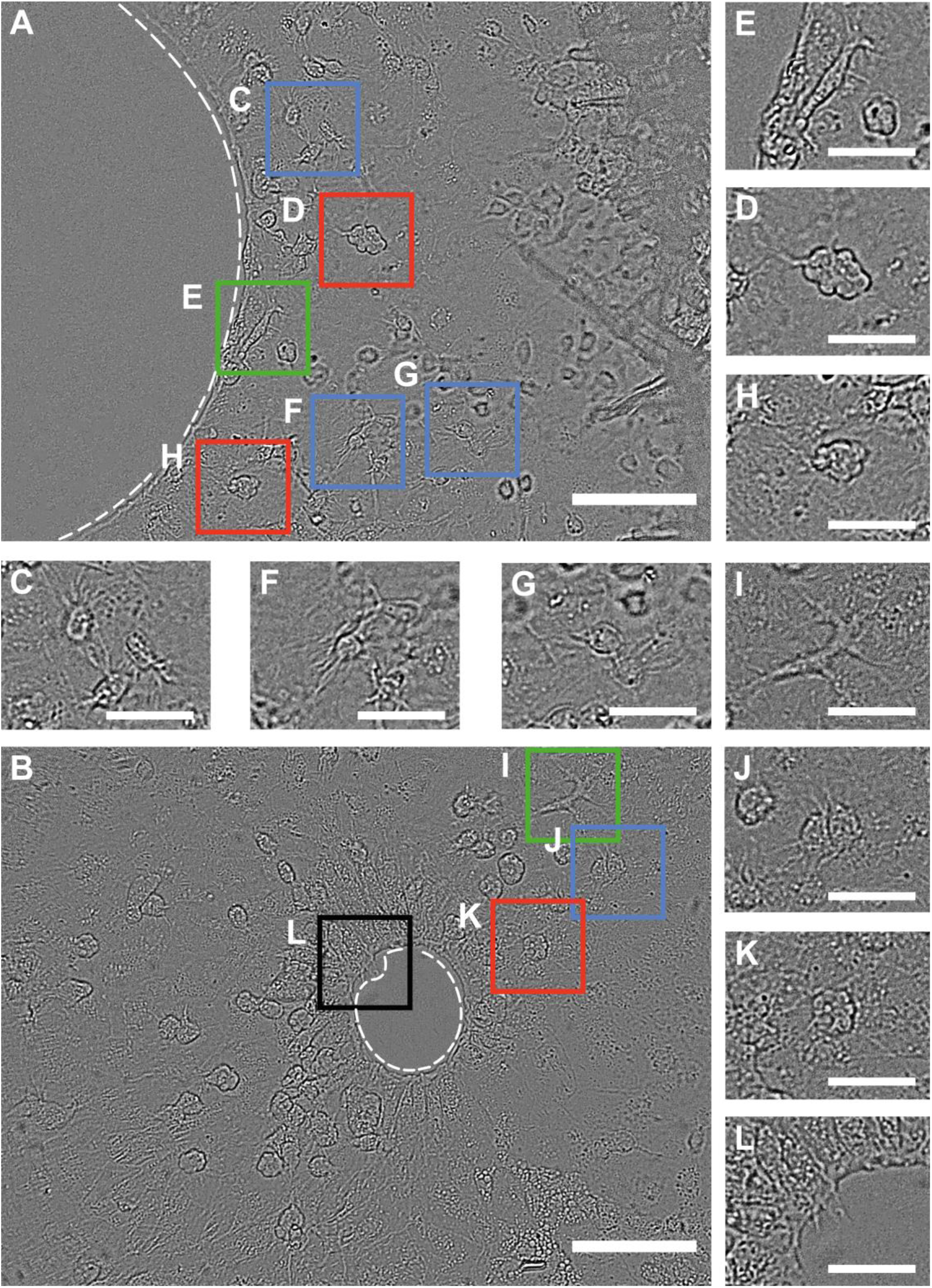
Mesohyl cells in the regenerative membrane (RM). (A-B) RM during closure, light microscopy; (C, F, G, J) – sclerocytes (blue frames); (D, H, K) – granular cells (red frames); (E, I) – large amoeboid cells (green frames); (L) – filopodia on the leading edge of the RM (black frame). Light microscopy, frames from time-lapse imaging. White dotted lines mark the leading edge of the RM. Scale bars (A, B) – 50 μm; (C-L) – 20 μm.

First and the most abundant type is round cells, 5-7 μm in diameter with long (up to 2-4 cell diameters) and thin filopodia (~5-10 per cell) (Fig. 3C, F, G, J). We often detected these cells in association with spicules at the late stage of regeneration, so we assume they are sclerocytes freely migrating in the RM (Movie S1, S3).

The second type, granular cells, their cell surface looks like it continuously undergoes ‘blebbing’ (Fig. 3D, H, K). It should also be noted that the number of granular cells in the RM is significantly higher than in the intact body wall.

The latter type is represented by large amoeboid cells. The long branching extensions are visible quite clearly; these tend to migrate using these extensions (Fig. 3E, I). In the intact body wall, these cells lie just above the surface of the choanoderm.

It is not known for certain whether mesohyl cells perform any function in the RM formation process, except for sclerocytes synthesising spicules during RM transformation into intact body wall (Lavrov *et al*., 2018) at the late stage of regeneration (Movie S3). However, all these cells tend to concentrate on the leading edge of the RM (Fig. 3A, B).

Mesohyl cell migration dynamics does not differ statistically between different cell types, so here we present generalised data. The average cell speed during the regeneration is 3.279±0.0917 μm/min (n=68). The migration efficiency of mesohyl cells is 0.5242±0.0247 (n=70).

### Actin structures in cells of the intact body wall

Intact exopinacocytes are T-shaped (Fig. 4), their cell body is submerged into the mesohyl matrix and has a few filopodia linking exopinacocytes to each other (Fig. 4B). Each exopinacocyte also has a flat polygonal external part (Fig. 4A), a cytoplasmic plate, which is connected with the cell body (Fig. 4B) through a thin ‘neck’. Cytoplasmic plates represent the outer layer of the body wall; thin actin bundles at their periphery delineate cell margins (Fig. 4A).

**Figure 4.**
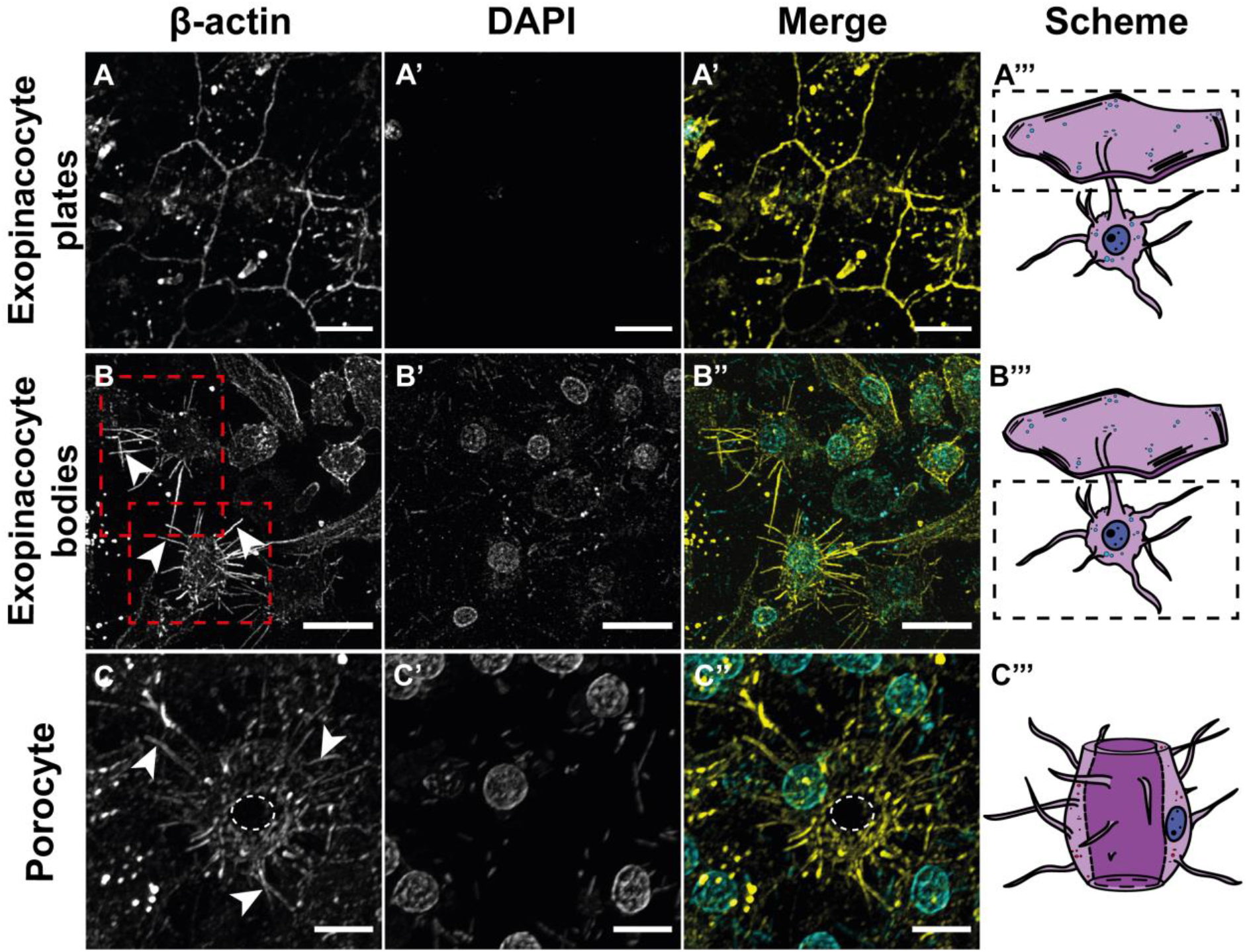
Actin filaments in the exopinacocytes and porocytes of the intact body wall. (A) – exopinacocyte cytoplasmic plates covering the body surface; (B) – exopinacocyte bodies submerged into mesohyl (red dotted lines); (C) – porocyte, the white dotted line marks ostia; CLSM (maximum intensity projection of several focal planes); cyan – DAPI, cell nuclei; yellow – cytoplasmic β-actin isoform, actin filaments. (A’’’), (B’’’), (C’’’) – cell schemes; the black dotted lines outline exopinacocytes parts demonstrated in the row; the white arrowheads mark filopodia. Scale bars: (A-B) – 10 μm; (C) – 5 μm.

Porocytes represent a specific cell type, penetrating the sponge body wall from the exopinacoderm to choanoderm (Fig. 4C). Porocyte is cylindrical cell with ostium – pass-through intracellular canal connecting the environment with the inner lumen of the sponge lined by the choanoderm. Actin filaments in porocytes are also represented by cortical actin bundles and numerous filopodia (Fig. 4C).

Choanocytes are tightly packed prismatic cells with an apical ring of actin-cored microvilli (Fig. 5A) located around a flagellum. Most actin structures in choanocytes are also cortical actin bundles (Fig. 5A-B).

**Figure 5.**
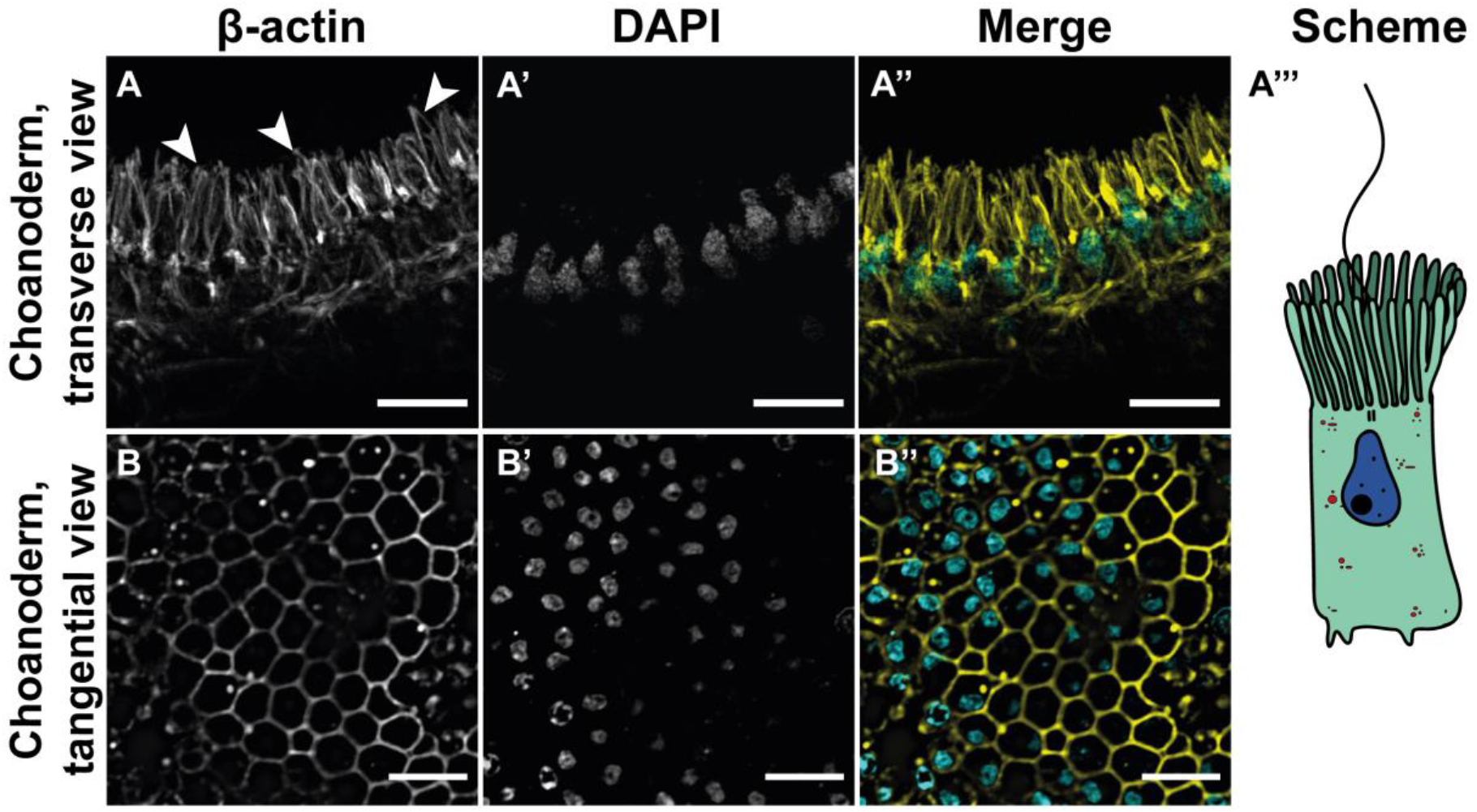
Actin filaments in the choanocytes of the intact body wall. (A) – choanoderm, transverse view; (B) – choanoderm, tangential view; CLSM (maximum intensity projection of several focal planes); cyan – DAPI, cell nuclei; yellow – phalloidin staining (A), cytoplasmic β-actin isoform (B), actin filaments. The white arrowheads mark microvilli collars. Scale bars (A-B) – 10 μm.

Mesohyl cells form a heterogeneous population. They differ in morphology and, most likely, in function. Using immunocytochemistry, we distinguished three cell types – large amoeboid cells, sclerocytes, and granular cells (Fig. 6).

**Figure 6.**
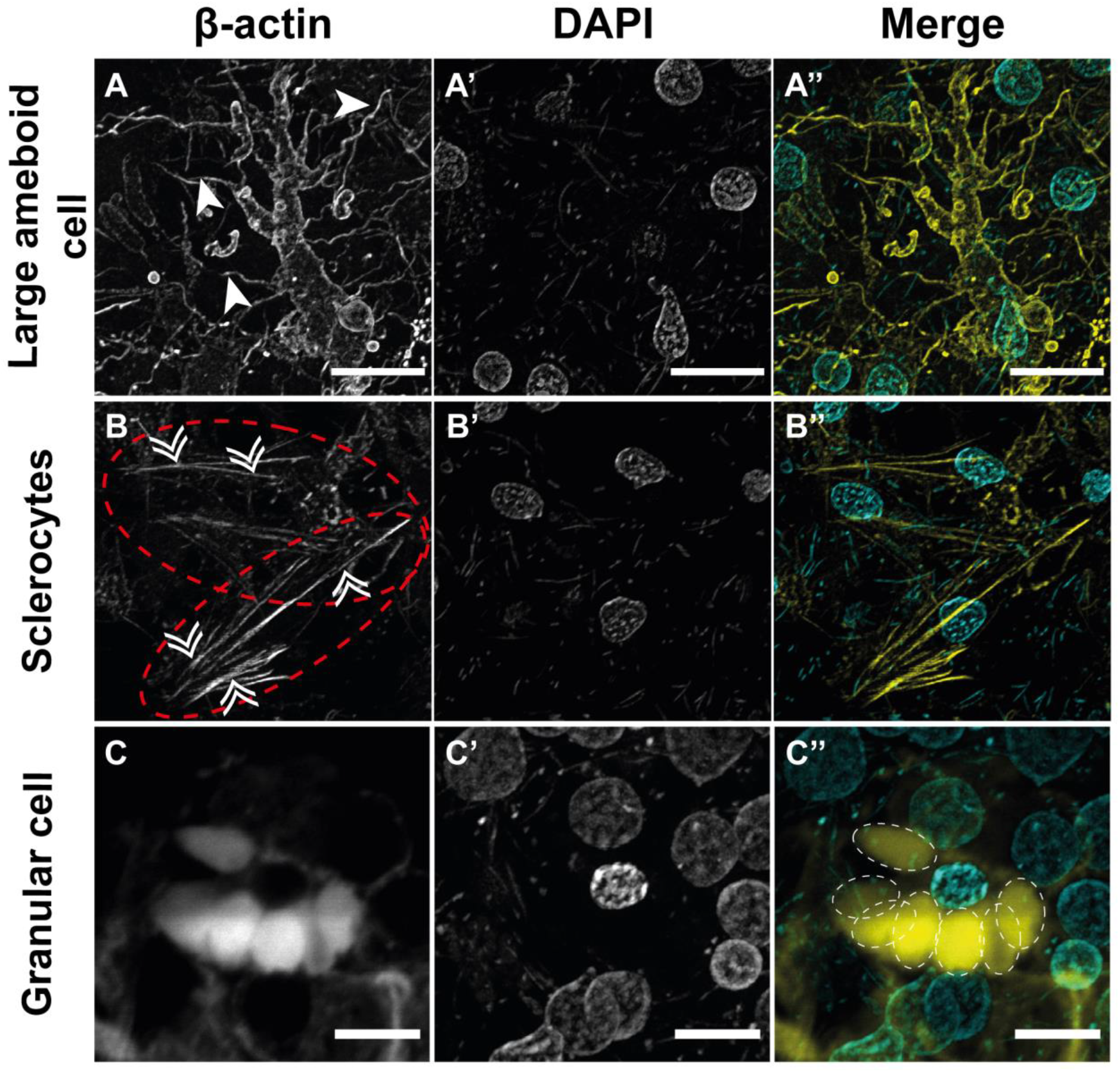
Actin filaments in the mesohyl cells of the intact body wall. (A) – large amoeboid cell; (B) – sclerocytes (red dotted lines); (C) – granular cell, the white dotted lines outline the edges of autofluorescent granules; CLSM (maximum intensity projection of several focal planes); cyan – DAPI, cell nuclei; yellow – cytoplasmic β-actin isoform, actin filaments. The white arrowheads mark filopodia, the double arrowheads mark stress fibers. Scale bars (A, B) – 10 μm; (C) – 5 μm.

Large amoeboid cells often have an oval nucleus and lobopodia getting thinner and branching at the end (Fig. 6A). Their typical location is in the mesohyl matrix, right above the choanoderm layer. The staining of these cells appears to be homogeneous probably because of background noise of the cytoplasm. Large amoeboid cells differ from other mesohyl cells due to their size (224.8±16.78 μm^2^, n=19) and lower circularity value (0.0816±0.0122, n=19) caused by branching extensions (Table S3). AR value is 1.837±0.09 (n=19) indicating that extensions are widely spread in numerous directions (Table S3).

Sclerocytes have a spindle-like shape, outlined with cortical actin, and often a few filopodia as well as numerous stress fibers along the major axis (Fig. 6B). In intact tissues, these cells have an intermediate area value – 113.9±4.275 μm^2^ (n=99) and a low circularity value 0.3048±0.0112 (n=99) (Table S3). Since the sclerocytes are elongated along with the axis of the spicule they synthesise, the AR value is quite high – 3.019±0.0933 (n=99) (Table S3).

Granular cells are more numerous in sponges, collected during autumn and winter. These cells are of intermediate size, with small condensed nucleus and cell edges aligned by cortical actin bundles. These cells have a lot of homogeneously stained large granules which alter the morphology of the cell due to dense packing (Fig. 6C). These cells also have an intermediate area values: 84.85±7.751 μm^2^ (n=6) and quite high circularity value – 0.6597±0,0501 (n=6) (Table S3). The AR value (1.555±0,0714 (n=6)) indicates the slightly elongated form of the cell (Table S3).

### Actin structures in cells of the regenerative membrane

The outer layer of the regenerative membrane is represented by exopinacocytes, and the inner layer, depending on the stage of regeneration, by endopinacocytes or redifferentiating choanocytes; with a thin layer of mesohyl containing single migrating cells (Fig. 7A).

**Figure 7.**
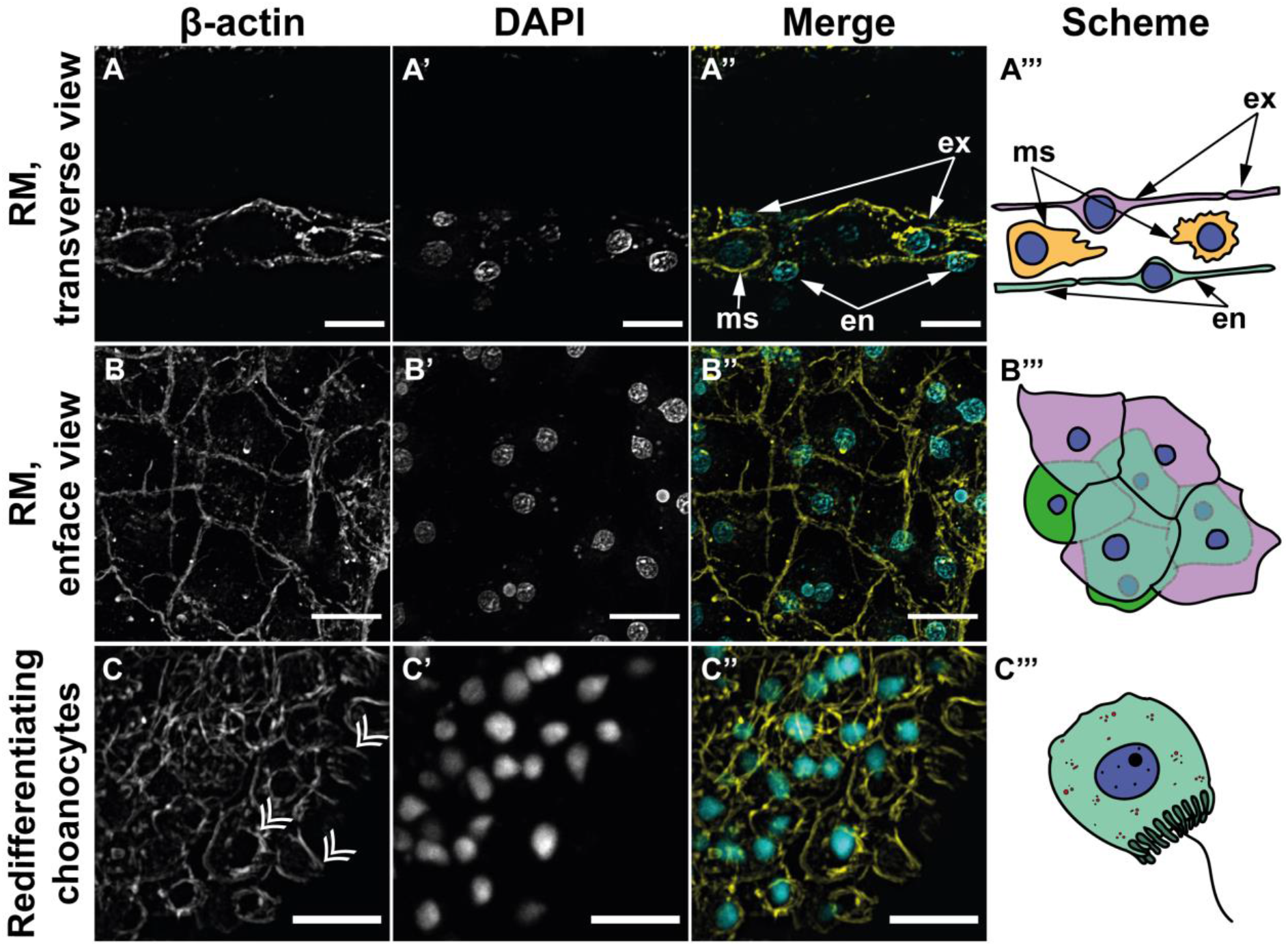
Actin filaments in the exopinacoderm, endopinacoderm, and choanoderm of the regenerative membrane (RM). (A) – RM, optical section demonstrating transverse view; (B) – RM, flat exo- and endopinacocytes; (C) – redifferentiating choanocytes at the periphery of RM; CLSM (maximum intensity projection of several focal planes); cyan – DAPI, cell nuclei; yellow – cytoplasmic β-actin isoform (A), phalloidin staining (B-C), actin filaments. (A’’’), (B’’’), (C’’’) – cells schemes. The double arrowheads mark choanocyte microvilli collar. en, endopinacocytes; ex, exopinacocyte; ms, mesohyl cell. Scale bars: (A-B) –15 μm; (C) – 10 μm.

The actin structures in flat polygonal exopinacocytes of RM are represented by cortical filament bundles (Fig. 7B). Pinacocytes on the leading edge of the RM tend to form thick actin cable (Fig. 8A). The cable runs continuously at the leading edge and is visualised until complete sealing of the RM. We have never observed similar actin structures in pinacocytes lying behind the leading edge. Cells at the leading edge of the wound also possess small short-living filopodia (lifetime is ~20 time frames), which are highly dynamic (Movie S1). Non-muscle myosin II colocalises along the cable in a beaded intermittent pattern (Fig. 8B). This indicates direct binding of myosin with the cable and allows us to define this structure as actomyosin cable with contractile properties.

**Figure 8.**
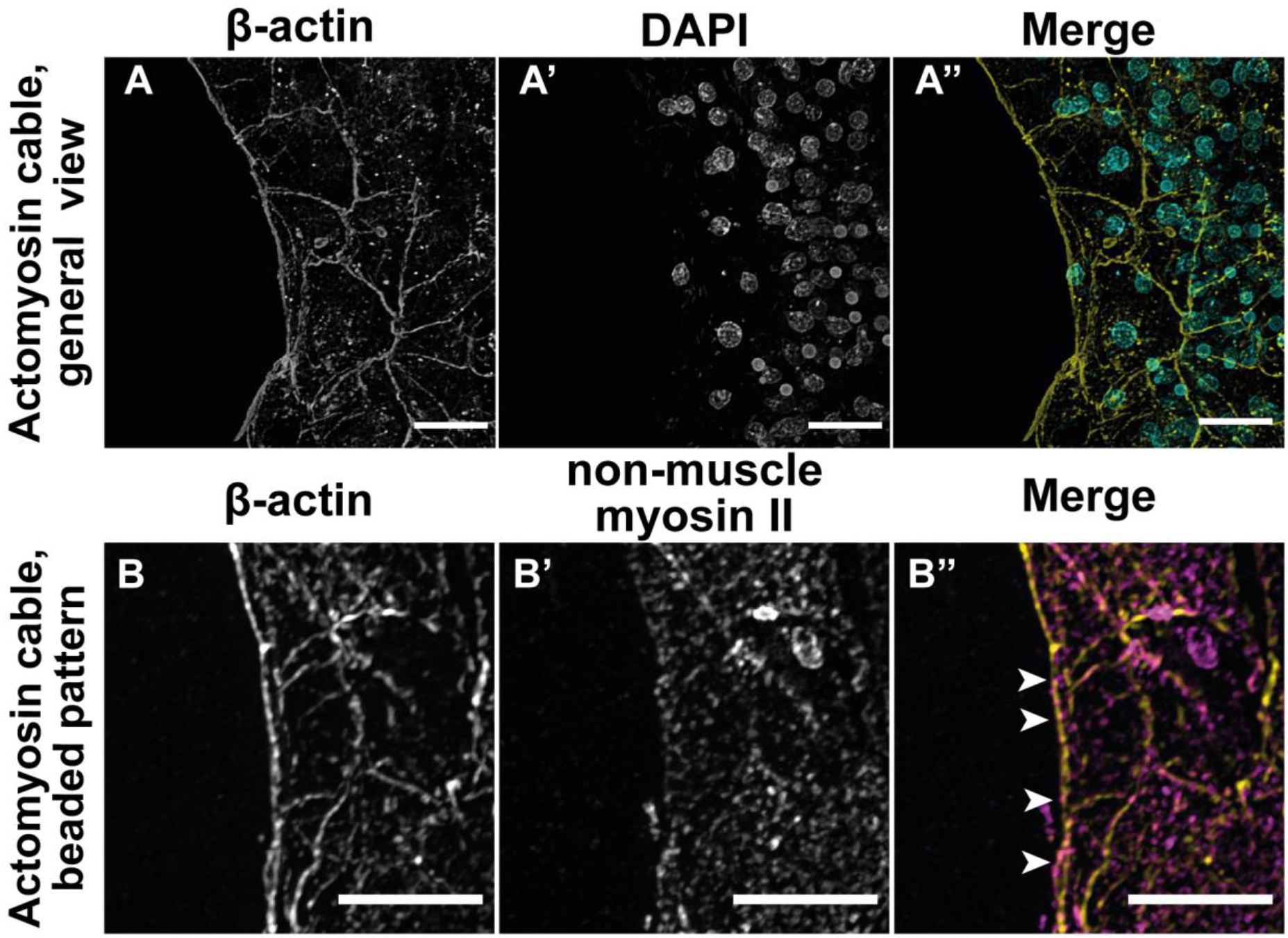
Actomyosin contractile cable at the leading edge of the regenerative membrane (RM). (A) – general view; (B) – beaded intermittent pattern of myosin/actin colocalisation; CLSM (maximum intensity projection of several focal planes); cyan – DAPI, cell nuclei; yellow – cytoplasmic β-actin isoform, actin filaments; magenta – non-muscle myosin II. The white arrowheads mark intermittent beaded pattern of myosin distribution. Scale bars (A) – 15 μm; (B) – 5 μm.

At the early stages of regeneration (0-48 hpo), the inner layer of the growing RM is represented by flat polygonal endopinacocytes (Fig. 7B). These cells do not have the microvilli collar and flagellum. The actin cytoskeleton in these cells consists of cortical bundles of filaments. The later stage of regeneration (48-96 hpo), when the transformation of the RM into the intact body wall occurs, is distinguished by the presence of ball-shaped cells with cortical actin filaments and short microvilli (Fig. 7C). These are redifferentiating choanocytes. They are packed loosely in comparison with tightly packed prismatic choanocytes of the intact body wall. Endopinacocytes redifferentiate into choanocytes gradually: from the peripheral sides of RM to its centre.

The mesohyl layer in the RM is quite thin; cells have actin protrusions and migrate intensively through the extracellular matrix. However, we have not seen any stress fibers in them.

### The morphological parameters of the cells change during regeneration

As RM growth is considered to be dependent on the transformations of epithelial-like cell layers (Korotkova, 1963; Lavrov *et al*., 2018; Caglar *et al*., 2021), we sought to estimate this process quantitatively. We describe the cell morphology in epithelial-like layers in the intact body wall and the RM, using three parameters: area, circularity and aspect ratio.

The intact choanocytes are small cells with an area of 29.74±0.515 μm^2^ (n=131) (Fig. 9A). These prismatic cells are almost round in the cross-section, so the circularity value is 0.907±0.004 (n=131) (Fig. 9B), and AR is 1.177±0.008 (n=129) (Fig. 9C).

**Figure 9.**
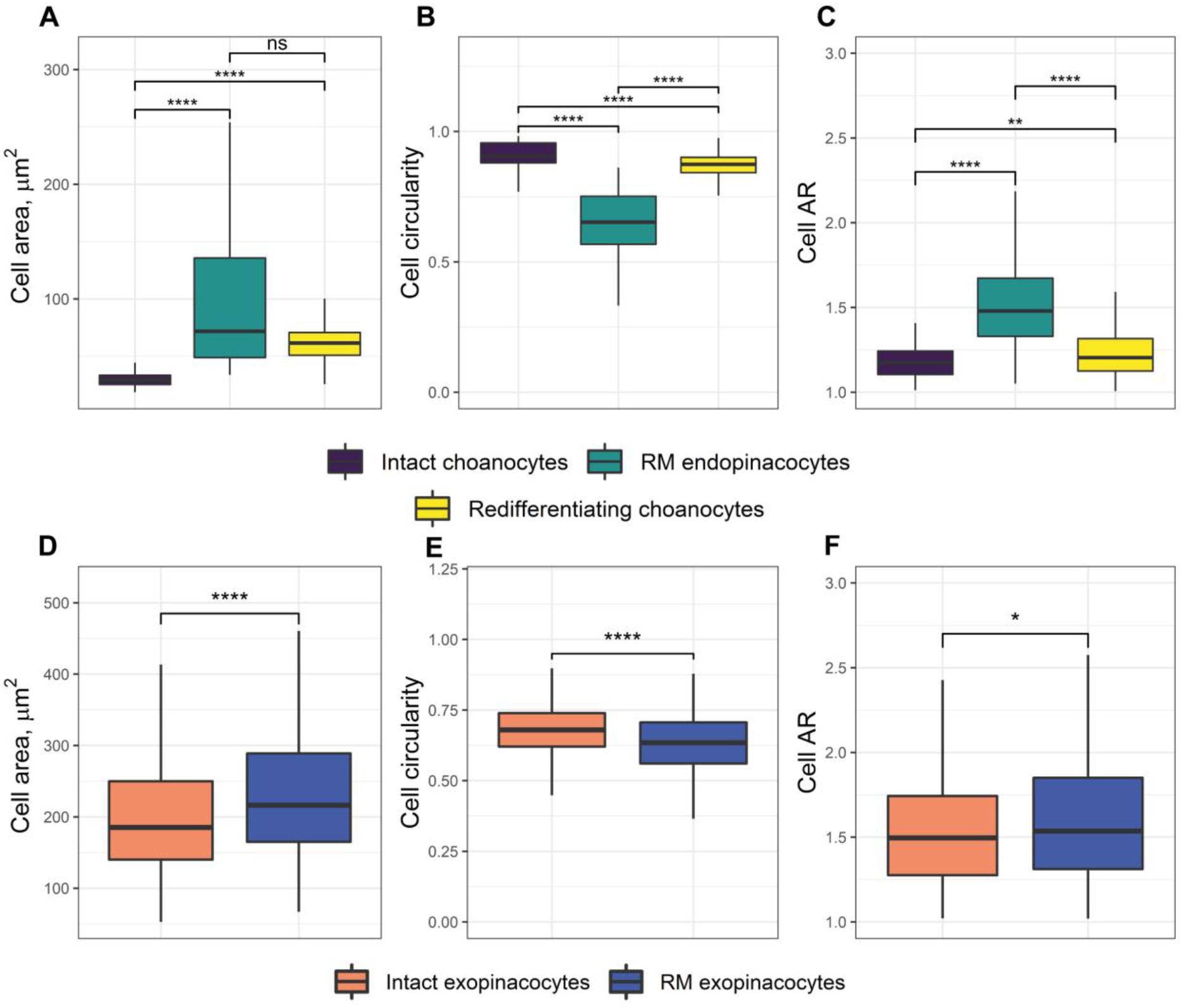
Morphometrical analysis of cell transformations in the choanoderm (A-C) and exopinacocyte (D-E) during regenerative membrane formation. (A, D) – cell area, (B, E) – circularity, (C, F) – aspect ratio. Data is shown with median values (thick horizontal lines), interquartile ranges (boxes) and total ranges (whiskers). Asterisks mark statistical significance between different cell types: p<0.0001 (****); p<0.001 (***); p<0.01(**); p<0.05 (*); ns – not significant. Dunn’s post hoc tests with the Benjamin-Hochberg adjustment (A-C); the one-sided Mann-Whitney U-test (D-E).Supplementary material

Transdifferentiation of choanocytes to endopinacocytes results in a significant area increase (Fig. 9A) – 89.63±7.535 μm^2^ (n=50, p-value=2.60e-31, Dunn’s multiple comparisons test with Benjamini & Hochberg correction, or ‘BH’), AR increase (Fig. 9C) up to 1.56±0.049 (n=51, p-value=8.09e-16, Dunn’s multiple comparisons test with BH), and circularity decrease (0.65±0.019, n=52, p-value=1.25e-32, Dunn’s multiple comparisons test with BH) (Fig. 9B).

Redifferentiating choanocytes at the late stages of regeneration have intermediate morphological parameters, which distinguish them from both intact choanocytes and membrane endopinacocytes. They have a rather high area value – 62.04±0.890 μm^2^ (n=285), significantly different (p-value=4.26e-54, Dunn’s multiple comparisons test with BH) from intact choanocytes (Fig. 9A). They also differ in the circularity value (0.881±0.002, n=245) (Fig. 9B) and AR (1.226±0.009, n=276) (Fig. 9C) both for choanocytes (p-value=1.03e-10 for circularity; p-value=4.54e-3 for AR, Dunn’s multiple comparison test with BH) and endopinacocytes (p-value=2.71e-17 for circularity; p-value=7.22e-12 for AR, Dunn’s multiple comparisons test with BH).

Intact exopinacocytes have large (199.6±5.188 μm^2^, n=266) (Fig. 9D) cytoplasmic plates with high circularity value (0.675±0.006, n=268) (Fig. 9E). The AR value of the plates (1.514±0.019, n=256) (Fig. 9F) indicates that the exopinacoderm is a stable cell layer and does not migrate in the intact body wall. The bodies of exopinacocytes merged into mesohyl has medium-size cell area (67.01±1.941 μm^2^, n=73) with low circularity rate (0.128±0.008, n=67) and low AR (1.547±0.037, n=67).

During the regeneration as the exopinacocytes flatten, the cell area significantly increases (p-value=9.97e-05, one-tailed Wilcoxon signed rank test) up to 231.2±6.367 μm^2^ (n=196) (Fig. 9D). Notably, the circularity of cells significantly decreases to 0.629±0.008 (Fig. 9E), (n=196, p-value=7.945e-06, one-tailed Wilcoxon signed rank test). The increase of AR value (1.587±0.027, n=185) is also statistically significant (p-value=0.03852, one-tailed Wilcoxon signed rank test) (Fig. 9F).

## Discussion

### The regeneration process in the calcareous sponge Leucosolenia variabilis

The regeneration in *L. variabilis* closely resembles this process in higher metazoans (Carlson, 2007), and could be split into generalised steps: wound healing, mobilisation of cell precursors, and morphogenesis (Tiozzo and Copley, 2015; Bideau, et al. 2021).

The isolation of the internal milieu during *L. variabilis* regenerations corresponds to the wound healing steps. It starts already at 1 hpo (Fig. S2) when leader cells of exopinacoderm spread and extend their filopodia with further contact with transdifferentiating choanocytes over the mesohyl layer (Lavrov *et al*., 2018). This process helps restoring the barrier against pathogens and prevent osmotic shock. These cell shape changes are known to be transcription-independent, which is why they appear quickly after wounding (Cordeiro and Jacinto, 2013).

The next step, the mobilisation of cell precursors, seems to be the key one, as it partially specifies the following morphogenesis type (Kawamura *et al*., 2008; Jopling *et al*., 2011; Gemberling *et al*., 2013). There are three main cell sources in regeneration: (1) proliferation and differentiation of adult stem cells; (2) dedifferentiation or transdifferentiation of somatic cells in an injured area; (3) proliferation of pre-existing fully differentiated somatic cells (Tiozzo and Copley, 2015). These sources are not self-exclusive and could coexist in various combinations in some species. The following morphogenesis leads to the reconstruction of lost body part, and could be epithelial or mesenchymal. The former type is based on transformations of epithelial cell layers in an injured area with minor or no contribution of cell proliferation. Mesenchymal morphogenesis, on the other hand, assumes blastema formation through intensive proliferation of mesenchymal cells undergoing the EMT and MET (Hay and Zuk, 1995; Vervoort, 2011; Borisenko *et al*., 2015; Alibardi, 2022; Tang *et al*., 2022).

In case of calcareous sponge regeneration, the mobilisation of cell precursors and following morphogenesis seem to be merged into a single phase: the formation of RM. It is the main and unique process during regeneration in calcareous sponge. RM growing occurs due to spreading of two epithelial-like cell layers, exopinacoderm and choanoderm, without any adhesive substrate or ECM matrix. There is neither blastema formation similar to the regeneration in Demospongiae (Borisenko *et al*., 2015; Ereskovsky *et al*., 2020), nor the increase in proliferation activity in *L. variabilis* tissues (Lavrov *et al*., 2018). Thus, the RM formation relies on epithelial morphogenesis and transformations of pre-existing cell layers. In this study, we aimed to trace cytoskeletal rearrangements underlying these morphological changes in the cells.

### The dynamics of regenerative membrane formation

The dynamics of RM formations has three key features: it is non-linear, stochastic (display growth fluctuations) and varies greatly among individuals. In the process of RM formation, there are periods when there is a much slower dynamics than in the main phase of growth. First of them is an initial lag period between end of wound healing (~1 hpo) and the first appearance of RM on the periphery of the wound, which occurs only after ~12 hpo. A similar lag period of 6 hours was observed during regeneration in the mouse corneal epithelium and was presumably associated with the reorganisation of cells at the leading edge (Danjo and Gipson, 1998). Another period of slower dynamics occurs at later stages of RM growth and explains the non-linear nature of curves in Fig. 2F. We associate it with a decrease in the perimeter of the wound and, accordingly, contact inhibition, which is triggered by the membrane cells coming into contact, similar to how it occurs in cellular monolayers (Lanosa and Colombo, 2008). We also assume that the filopodia that appear in the cells of the leading edge of the wound (exopinacocytes) might modulate this process similar to those involved in *Drosophila* dorsal closure. In that model, filopodia and lamellipodia on the opposite leading edges contact and draw them together (Jacinto *et al*., 2000; Garcia-Fernandez *et al*., 2009).

The observed fluctuations in the dynamics of RM growth can be explained by the mechanical stress to which the leading edge of RM is subjected: time-lapse imaging allows to visualise tears occurring at the edge indicating the mechanical tension applied to the RM. Such tears cause a local retraction of the leading edge and, thus, temporary increase in the area of the wound. Similar observations were obtained in MDCK cells in a model of non-adhesive epithelialization, where the wound in the monolayer was tightened by a stochastic sequence of contractions and increases in its area, and the observed amplitudes were up to 1/5 of the entire wound size (Nier *et al*., 2015).

We assume that the heterogeneity of dynamics among individual specimens is caused by different physiological states, the initial area of the wound, the conditions of the environment from which the animal was taken (atmospheric pressure, the strength of tidal influences, water temperature, season). It is known, for example, that the physiological state of sponges is affected by the temperature of the environment, its significant increase results in the mass death of individuals (Ereskovsky *et al*., 2019, 2021; Lavrov *et al*., 2022). The physiological state of an individual in turn directly affects the course of various restorative processes in sponges (Valisano *et al*., 2006; Lavrov *et al*., 2020; Ereskovsky *et al*., 2021).

### Mesohyl cells: morphology and contribution to regeneration

Unlike pinacoderm and choanoderm, mesohyl is structureless, its cells constantly moving and do not form any stable conglomerates. Their morphology and possible contribution to regeneration will be discussed in this section.

Mesohyl cells vary in morphology and perform multifarious functions. The role of spicule-synthesising cells, sclerocytes, seems to be the clearest and studied one. These cells in calcareous sponges have well-established septate junctions (Ledger, 1975; Lavrov et al., 2018) which probably perform a barrier function, maintaining the ionic gradient between the extracellular spicule cavity and the mesohyl matrix. Thus, it is consistent that sclerocytes have thick actin bundles outlining cell edges since there is evidence septate junctions are associated with actin filaments (Lane and Flores, 1988).

Granular cells have season-dependent distribution in *L. variabilis* tissues: these cells are absent in spring and early summer samples, but autumn and winter samples have plenty of them. The reason for the homogeneous staining of the granular component is not clear and requires further research using other methods and dyes. The exact function of granular cells is not established but can be attributed to storage functions (store various metabolites or symbiotic bacteria), antimicrobial activity or reproduction process (Simpson, 1984; Gauthier, 2009; Ereskovsky and Lavrov, 2021).

Large amoeboid cells remain an enigma. The most morphologically similar cell type is fiber cells in *Trichoplax adhaerens* which also have numerous long branching extensions. In Placozoa these cells connect the ventral and dorsal epithelia to each other and their function remains unclear (Smith *et al*., 2014). Actin filaments in fiber cells form a dense meshwork which probably provides their ability to contract (Behrendt *et al*., 1986; Thiemann *et al*., 1989). They also have large autofluorescent granules, and therefore their staining appears homogeneous (Smith and Reese, 2016). Large amoeboid cells of *L. variabilis* always lie above the choanoderm, thus, we assume they provide some kind of support for choanocytes either signal or mechanical. Or *vice versa*, large amoeboid cells transmit signals and (or) nutritional chemicals from the choanoderm to other cells. However, their function requires further research.

Porocytes, tubular cells through which water penetrates from the external environment to the spongocoel, are not considered to be mesohyl cells, but will be considered here as not directly involved in the formation of RM. Porocytes of calcareous sponges are capable of contracting and, therefore, regulating water flow through the animal body (Jones, 1966; Eerkes-Medrano and Leys, 2006). The actin cytoskeleton of porocytes forms a widely spaced grid and may fit to constant contractions. This assumption seems especially relevant due to the establishment of contractile actin bundles that contain striated muscle myosin II and are regulated by myosin-light-chain kinase (MLCK) in the sponge *Ephydatia muelleri* tissues (Colgren and Nichols, 2022). Numerous filopodia of porocytes are similar to those in exopinacocytes bodies have and will be discussed later. During regeneration, porocytes appear synchronously with redifferentiation of endopinacocytes into choanocytes, from the periphery of the RM to its centre. In this work, we did not track the source of these cells, but in an earlier study it was shown that porocytes are the result of the transdifferentiation of RM exopinacocytes (Lavrov *et al*., 2018).

At the moment, the role of mesohyl cells in the RM formation has not been clarified. Ultrastructural studies of the regeneration of calcareous sponges have not revealed the contribution of mesohyl cells: they do not integrate into the RM, do not participate in its formation (Lavrov *et al*., 2018; Ereskovsky *et al*., 2021). However, our time-lapse imaging illustrates that mesohyl cells tend to concentrate at the leading edge of the RM. In addition to sclerocytes, we detect two more types of cells in the membrane: large amoeboid cells and granular cells.

The number of granular cells in RM significantly exceeds their number in intact tissues. This might be an argument for their possible role in the formation and transformation of RM. Granular cells might transport nutrients necessary for the body wall formation or substances with antimicrobial activity preventing exceed development of microorganisms at the wound site. Alternatively, granular cells might accumulate metabolites that have appeared as a result of a violation of the integrity of tissues.

We also assume that mesohyl cells could accumulate at the leading edge of the RM to intensify ECM synthesis in the forming tissue, since it is their main function as resident in this analogue of the connective tissue of Eumetazoans (Ereskovsky, 2010).

### RM formation as a result of epithelial morphogenetic processes

The pinacoderm in asconoid sponges forms the outer cover of the body (exopinacoderm) and outlines the inner part of the oscular rim (endopinacoderm) (Bergquist, 2001; Eerkes-Medrano and Leys, 2006; Ereskovsky and Lavrov, 2021). In *L. variabilis*, intact exopinacocytes are T-shaped. Such type of organisation (called ‘insunk epithelium’) is also present in many Demospongiae species (Simpson, 1984) and in some species from other phyla: epithelium of *Convoluta convoluta* (Xenacoelomorpha) (Pedersen, 1964), epithelium of some lecithoepitheliates and bdellourid triclads (Tyler and Hooge, 2004) and dorsal epithelial cells in *Trichoplax adhaerens* (Placozoa) (Smith *et al*., 2014). Therefore, such morphology is not unique and is probably associated with a certain plasticity of the epithelial tissues.

And yet intact *L. variabilis* exopinacocytes have a unique feature, reported for the first time in our work: filopodial extensions connecting cell bodies with each other. We are not well aware what function these extensions fulfil, still, there are some assumptions. First, they might provide mechanical integrity and stability of the cell layer as there is no evidence of conventional cell-cell junctions between cytoplasmic plates, though there are many studies confirming the presence of cell adhesion machinery in sponges (Boury-Esnault *et al*., 2003; Adell *et al*., 2004; Leys *et al*., 2009; Leys and Hill, 2012; Leys and Riesgo, 2012; Jonusaite *et al*., 2016; Belahbib *et al*., 2018; Mitchell and Nichols, 2019). Second assumption is that extensions connecting exopinacocytes provide signalling and temporally and spatially coordinated contraction of exopinacoderm and, thus, regulate water flow through the body, similar to previously observed (Jones, 1957; Elliott and Leys, 2007; Nickel *et al*., 2011).

The outer layer of RM consists of flat polygonal exopinacocytes. These cells represent a result of fusion between cytoplasmic plate and cell body accompanied by substantial changes in actin cytoskeleton structure. In marginal exopinacocytes of RM cortical actin bundles are replaced by filopodia on the wound side and remain stable at the other sides of the cells. These either mechanosensing or signalling protruding structures probably perform a similar function as was shown in *Drosophila* dorsal closure – sealing the opposite edges of epithelium (Stramer *et al*., 2005; Garcia-Fernandez *et al*., 2009; Abreu-Blanco *et al*., 2012). The same cells (RM exopinacocytes) form an actomyosin cable on the edges of the wound.

Our morphometric analysis indicates the presence of polarisation in the epithelial-like cell layer of exopinacoderm during RM formation: circularity decreases and AR increases. However, there are no common or predicted migratory structures (lamellipodia, filopodia or lobopodia) and no other features of collective cell crawling detected. During flattening, exopinacocytes do not lose intercellular junctions, they move collectively, though there are no leader cells typical for the collective cell migration mechanism of epithelization (Reffay *et al*., 2014). We also did not observe cryptic lamellipodia in cells of the second, third and more distant rows from the leading edge, although it has been shown that such structures might contribute to collective cell migration (Farooqui and Fenteany, 2005). We assume that the decrease in circularity is caused by the appearance of filopodia and the cell elongation toward the wound center. This as well explains the observed AR changes. We also associate cell elongation with mechanical tension generated by actomyosin cable contraction, as previously shown in cell monolayer epithelization (Lee and Gotlieb, 2005; Anon *et al*., 2012) and *Drosophila* embryogenesis (Jankovics and Brunner, 2006; Garcia-Fernandez *et al*., 2009). Hereby, morphological changes of individual exopinacocytes indicate fundamental transformations of the epithelial-like cell layer of the exopinacoderm.

The choanoderm in the asconoid body of *L. variabilis* represents the inner layer of cells. It provides feeding, excretion, oxygenation, and gametogenesis (Simpson, 1984). Choanocytes are polarised prismatic cells with a flagellum and a microvilli collar on the apical pole and cytoplasmic extensions anchoring the cell to the mesohyl matrix at the basolateral domain (Ereskovsky and Lavrov, 2021). The general appearance of choanoderm in Calcarea and Demospongiae includes clear morphological apicobasal polarity, structures similar to cell junctions between choanocytes (Eerkes-Medrano and Leys, 2006; Lavrov *et al*., 2018, 2022), numerous cellular junction orthologous proteins (Leys and Hill, 2012; Peña *et al*., 2016; Miller *et al*., 2018; Mitchell and Nichols, 2019) as well as basal lamina components (Boute *et al*., 1996; Fahey and Degnan, 2010; Leys and Riesgo, 2012) providing strong evidence for claiming choanoderm as epithelial-like cell layer. This assertion is confirmed by subcellular cytoskeletal organisation of choanocytes provided in this study. The actin cytoskeleton in choanocytes is quite similar to those reported in prismatic epithelia of Eumetazoa (Bretscher and Weber, 1978; Kukulies *et al*., 1984; Waterman-Storer *et al*., 2000) – actin bundles act as microvilli core and are oriented along the apicobasal axis of the cells.

It is generally accepted to call transformations of sponge cells transdifferentiation since cells completely change their morphological appearance and functional role (Borisenko *et al*., 2015; Ereskovsky *et al*., 2015; Adamska, 2018; Lavrov *et al*., 2018; Caglar *et al*., 2021). Therefore, transformation of choanocytes with loss of some morphological features (microvilli collar and flagellum) following by the loss of filtering ability can also be considered as a transdifferentiation.

The flattening of originally columnar choanocytes and, respectively, their area increase, circularity decrease and AR increase, provide a source (together with exopinacocytes) for covering the excised area. Similar morphological changes are observed during intestinal epithelial restitution (Tétreault *et al*., 2005). Furthermore, such strong changes are only possible due to fundamental rearrangements of the cytoskeleton. For example, during *Drosophila* development, cells of amnioserosa flatten end elongate autonomously by so-called ‘rotary cell elongation’ (Pope and Harris, 2008). This mechanism relies on coordinated reorganisation of both microtubule and actin filaments without cell contact disruption. Transdifferentiation of choanocytes probably follows the same route considering the observed morphological changes: cell area and AR increase and circularity decrease. At the same time, in the endopinacoderm, as well as in the exopinacoderm, we did not observe structures responsible for the migration of cell layer as well as any leader cells.

The reverse process, columnarisation of flat endopinacocytes and restoration of choanocyte morphological characteristics, is referred here as redifferentiation as in previous studies (Lavrov *et al*., 2018; Ereskovsky *et al*., 2020). During redifferentiation endopinacocytes become spherical and, correspondingly, cell area and AR decrease, while circularity increases. Later stages are accompanied by further apicobasal elongation of cells (Lavrov *et al*., 2018). This process resembles terminal differentiation of kidney epithelia, when (1) cells establish terminal web and brush-border microvilli on the apical side, (2) their nuclei move to a basal cell side, and (3) cells acquire a columnar shape (Vijayakumar *et al*., 1999). There is no evidence of the existence of a terminal web in choanocytes, probably due to poor actin preservation during the preparation of TEM samples. However, other listed morphological features specific to these cells are present and become evident during the transformation of RM into the body wall.

Interestingly, the microvilli collar and the flagellum are the last to be lost during transdifferentiation and the first to re-establish during redifferentiation. This might be important because of the main function of choanocytes – water filtration – and capability of partially performing this function even with altered filtration apparatus.

Cell transformations during RM formation are fundamentally similar to those characteristics of higher animals, at least at the level of cell morphology and actin filament rearrangements. Thus, we assume that regeneration in *L. variabilis* is driven by epithelial morphogenesis with some distinct features: the absence of a substrate for cell movement and the presence of actin cable at the leading edge of the RM.

### RM formation as an analogue of epithelization over a non-adhesive substrate

Epithelial healing is based on two interacting mechanisms: collective cell crawling and actin ‘purse-string’ contraction. It was widely accepted that the first mechanism is more peculiar for adult animals (Zahm *et al*., 1991; Nusrat *et al*., 1992; Zahm *et al*., 1997; Yin *et al*., 2008) and the second one is typical for embryonic wounds and development (Martin and Lewis, 1992; Brock *et al*., 1996; Jacinto *et al*., 2000). Yet, at the moment, the number of works indicating these mechanisms work conjointly is increasing. Some researchers detect these mechanisms at different stages of the wound healing process (Brugués *et al*., 2014; Kuipers *et al*., 2014), others find them interacting throughout the time of healing process (Fenteany *et al*., 2000; Klarlund, 2012; Richardson *et al*., 2016; Kamran *et al*., 2017). There are also studies debating if the mechanism is determined by the size and (or) geometry of the wound (Bement *et al*., 1993; Danjo and Gipson, 1998; Grasso *et al*., 2007).

In the last few years, the model of “non-adhesive epithelization” has arise. This model assumes the absence of an adhesive substrate at the site of a wound: cells are not allowed to spread, flatten and migrate over the wound and therefore form so-called ‘epithelial bridges’ stretched over a non-adhesive substrate (Vedula *et al*., 2014; Albert and Schwarz, 2016). Their formation relies on the functioning of the actomyosin cable in cells from the ‘bridge’, connected not only to each other, but also to cells on an adhesive substrate around the wound site. The whole structure is also subjected to significant mechanical tension transmitted between neighbouring cells (Nier *et al*., 2015; Chen *et al*., 2019).

As was already mentioned, the RM formation occurs without a substrate, the membrane encloses over a gap in the sponge body wall. The morphology of RM is quite similar to the ‘epithelial bridges’ obtained by Vedula and co-authors (2014) during cultivating the MDCK line on fibronectin strips separated by a non-adhesive substrate. In that study, the formation of ‘bridges’ was slowed down if the distance between the fibronectin strips was increased to ~200 μm. However, our observations show that the formation of an actomyosin cable allows RM to overcome such distances with no issues and a wound with an initial radius of ~ 1000 μm can be rapidly sealed.

It is also known that epithelialization through the mechanism of actomyosin ring contraction requires the presence of cell-cell junction proteins which form clusters along the leading edge: cadherins (Brock *et al*., 1996; Danjo and Gipson, 1998; Brugués *et al*., 2014; Vedula *et al*., 2014), ZO-1 (Bement *et al*., 1993; Brugués *et al*., 2014), vinculin (Grasso *et al*., 2007). Some of them affect actin cable stability: for example, monoclonal anti-E-cadherin antibodies (ECCD-1) disrupt wound healing by the purse-string mechanism in mouse corneal epithelium (Danjo and Gipson, 1998). This additionally convinces us that there should be cell junctions between sponge cells that resemble those belonging to Eumetazoans, but they may differ in morphology and protein composition. In consistence with this evidence there are studies indicating orthologous cell-cell junction proteins in sponge tissues (Boury-Esnault *et al*., 2003; Adell *et al*., 2004; Eerkes-Medrano and Leys, 2006; Leys and Riesgo, 2012; Miller *et al*., 2018). The presence of cell junctions is driven by the necessity of stabilisation of the actin cable, subjected to significant mechanical tension. Disruptions and subsequent retractions of the RM leading edge, in addition to the already mentioned tension, might be a consequence of the force generated by the water flow through the body fragments. That is an additional obstacle to stabilising the actin cable.

The initial wound size and characteristic asymmetry of RM growth suggest we observe not a complete actomyosin cable formed along the entire perimeter of the wound, but shorter fragments (‘actin arcs’) as is typical for the epithelization of large wounds (Bement *et al*., 1993). Each of these fragments generates mechanical forces in its own sector of the RM.

Summing up, we assume that the formation of a RM during the regeneration in *L. variabilis* depends mainly on the contractile activity of the actomyosin cable, which coordinates extension of the cell layers of the exopinacoderm and endopinacoderm (transdifferentiated choanoderm) to the centre of the wound, and ensures the direction and constancy of their movement in non-adhesive substrate conditions. However, the question about the origin of force pushing the leading edge of the cell layer toward the centre of the wound remains relevant (Fenteany *et al*., 2000; Anon *et al*., 2012). Some studies indicate the cell crawling as the main way for wound closure while actin cable functions as a guiding structure (Anon *et al*., 2012). Others suggest that actomyosin cable assembly and contraction draw adjacent edges of the wound together (Tamada *et al*., 2007; Vedula *et al*., 2015).

## Conclusions

Regenerative membrane formation during regeneration in calcareous sponges is dependent on exopinacoderm and choanoderm coordinated flattening accompanied by the choanocyte transdifferentiation with temporary loss of the morphological features and physiological function. This process proceeds without a contribution of proliferation and, at some points, resembles ‘non-adhesive epithelization’: it has similar dynamics, occurs without substrate, and probably depend on the actomyosin cable formation. Morphological changes of the cells and alterations in their actin cytoskeleton are similar to those characteristic to higher Metazoans and, thus, are conservative through evolution.

Our data provide a basis for a further mechanistic analysis of morphogenesis in sponges. For example, the regulation of cytoskeleton components (actin filaments, microtubules) in sponges remains poorly characterised. This is also true for interactions and mutual regulation of microtubule systems and actin filaments, their role in morphogenetic processes. Forces driving RM formation and sealing also remain under elucidated. The role of Rho family GTPases and their effector proteins as well as microtubules and their associated proteins in regeneration of calcareous sponges are promising areas for future studies. Investigations in this direction will be an important step towards describing the mechanisms of morphogenetic processes in the basal groups of multicellular animals and the role of the cytoskeleton in these processes, which is the basis for identifying the mechanisms establishing true tissues in evolution.

## List of abbreviations

AJ: adherens junctions
AR: aspect ratio
CLSM: confocal laser scanning microscopy
DAPI: 4’,6-diamidino-2-phenylindole
ECM: extracellular matrix
EMT: epithelial-mesenchymal transition
FSW: filtered seawater
GTPase: nucleotide guanosine triphosphate (GTP) hydrolase
hpo: hours post operation
MET: mesenchymal-epithelial transition
MLCK: myosin light-chain kinase
RM: regenerative membrane
TEM: transmission electron microscopy
PBS: phosphate-buffered saline

## Acknowledgements

The authors acknowledge the support of Lomonosov Moscow State University Program of Development (Nikon A1 CLSM) and Centre of microscopy WSBS MSU. Authors sincerely thank Daniyal Saidov (Lomonosov Moscow State University) for statistical analysis tips, Elena Voronezhskaya (Institute of Developmental Biology, Russian Academy of Sciences) for fixation recommendations, Nikolay Melnikov, Anastasiia Kovaleva, Anna Tvorogova, Stani slav Kremnyov (Lomonosov Moscow State University) and Alexander Ereskovsky (Institute of Developmental Biology, Russian Academy of Sciences) for helpful tips and advice.

## Author contributions

KS, AL and AS designed the study. KS, AL and FB collected the material, carried out CLSM cytoskeleton studies, analysed and visualised data. KS conducted experiments and performed statistical analysis and its visualisation as well as time-lapse imaging and its post-processing. AL, KS and AS prepared the manuscript with contributions from all authors. All authors reviewed and approved the final manuscript.

## Data availability statement

Raw data were generated at Lomonosov Moscow State University, Biological faculty. Derived data supporting the findings of this study are available from the corresponding author Kseniia V. Skorentseva on request.

## Supplementary material

1. **Table S1.** Morphometric cell parameters (area, circularity, aspect ratio), raw data, epithelial-like cell layers.
2. **Table S2.** Morphometric cell parameters (area, circularity, aspect ratio), raw data, mesohyl cells.
3. **Table S3.** Descriptive statistics of morphometric cell parameters.
4. **Figure S1.** Scheme illustrating experimental procedures. FSW - filtered sea water; hpo – hours post operation; RM – regenerative membrane.
5. **Figure S2.** Wound edge, 1 hpo. Black arrows point towards transdifferentiating choanocytes of the inner side of the body wall. Scanning electron microscopy. Sample preparation described in Lavrov *et al*., 2018.
6. **Movie S1.** Regenerative membrane growth and sealing (part 1); exopinacocytes trembling and mesohyl cells migration in a fully sealed regenerative membrane (part 2). Time-lapse imaging. Video runs at 225-235x real time. Red arrows indicate filopodial activity at the leading edge of RM. Scale bar 50 μm.
7. **Movie S2.** Mechanical tension causes RM tearing and leading edge retraction. Time-lapse imaging. Video runs at 295x real time. Red arrows indicate rupture site at the leading edge of RM. Scale bar 50 μm.
8. **Movie S3.** Sclerocytes synthesising spicules in the regenerative membrane. Time-lapse imaging. Video runs at 225x real time. Scale bar 50 μm.

## References

Abreu-Blanco, M. T. et al. (2012) ‘Drosophila embryos close epithelial wounds using a combination of cellular protrusions and an actomyosin purse string’, Journal of Cell Science, 125(24), pp. 5984–5997. doi: 10.1242/jcs.109066.

Adamska, M. (2018) ‘Differentiation and Transdifferentiation of Sponge Cells’, Results and Problems in Cell Differentiation, 65, pp. 229–253. doi: 10.1007/978-3-319-92486-1_12.

Adell, T. et al. (2004) ‘Evolution of metazoan cell junction proteins: The scaffold protein MAGI and the transmembrane receptor tetraspanin in the demosponge Suberites domuncula’, Journal of Molecular Evolution, 59(1), pp. 41–50. doi: 10.1007/s00239-004-2602-2.

Albert, P. J. and Schwarz, U. S. (2016) ‘Dynamics of Cell Ensembles on Adhesive Micropatterns: Bridging the Gap between Single Cell Spreading and Collective Cell Migration’, PLoS Computational Biology, 12(4), pp. 1–34. doi: 10.1371/journal.pcbi.1004863.

Alexander, B. E. et al. (2015) ‘Cell kinetics during regeneration in the sponge *Halisarca caerulea:* How local is the response to tissue damage?’, PeerJ, 2015(3), pp. 1–19. doi: 10.7717/peerj.820.

Alibardi, L. (2022) ‘Activation of cell adhesion molecules and Snail during epithelial to mesenchymal transition prior to formation of the regenerative tail blastema in lizards’, Journal of experimental zoology. Part B, Molecular and developmental evolution, doi: 10.1002/JEZ.B.23139.

Anon, E. et al. (2012) ‘Cell crawling mediates collective cell migration to close undamaged epithelial gaps’, Proceedings of the National Academy of Sciences of the United States of America, 109(27), pp. 10891–10896. doi: 10.1073/pnas.1117814109.

Babbin, B. A. et al. (2009) ‘Non-muscle myosin IIA differentially regulates intestinal epithelial cell restitution and matrix invasion’, American Journal of Pathology, 174(2), pp. 436–448. doi: 10.2353/ajpath.2009.080171.

Behrendt, G. et al. (1986) ‘The ventral epithelium of *Trichoplax adhaerens* (Placozoa): Cytoskeletal structures, teil contacts and endocytosis’, Zoomorphology, 200, pp. 123–130.

Belahbib, H. et al. (2018) ‘New genomic data and analyses challenge the traditional vision of animal epithelium evolution’, BMC Genomics, 19(1), pp. 1–15. doi: 10.1186/s12864-018-4715-9.

Bement, W. M., Forscher, P. and Mooseker, M. S. (1993) ‘A novel cytoskeletal structure involved in purse string wound closure and cell polarity maintenance’, Journal of Cell Biology, 121(3), pp. 565–578. doi: 10.1083/jcb.121.3.565.

Bergquist, P. R. (1978) Sponges, University of California Press.

Bideau, L. et al. (2021) ‘Animal regeneration in the era of transcriptomics’, Cellular and Molecular Life Sciences, 78(8), pp. 3941–3956. doi: 10.1007/s00018-021-03760-7.

Blanchoin, L. et al. (2015) ‘Dynamic reorganization of the actin cytoskeleton’, F1000Research, 4(0), pp. 1–11. doi: 10.12688/f1000research.6374.1.

Borisenko, I. E. et al. (2015) ‘Transdifferentiation is a driving force of regeneration in *Halisarca dujardini* (Demospongiae, Porifera)’, PeerJ, 3(8), p. e1211. doi: 10.7717/peerj.1211.

Boury-Esnault, N. et al. (2003) ‘Larval development in the Homoscleromorpha’, Invertebrate Biology, 122(3), pp. 187–202.

Boute, N. et al. (1996) ‘Type IV collagen in sponges, the missing link in basement membrane ubiquity’, Biology of the cell, 88(1–2), pp. 37–44. doi: 10.1016/S0248-4900(97)86829-3.

Bretscher, A. and Weber, K. (1978) ‘Localization of actin and microfilament-associated proteins in the microvilli and terminal web of the intestinal brush border by immunofluorescence microscopy’, The Journal of cell biology, 79(3), pp. 839–845. doi: 10.1083/JCB.79.3.839.

Brock, J. et al. (1996) ‘Healing of incisional wounds in the embryonic chick wing bud: Characterization of the actin purse-string and demonstration of a requirement for Rho activation’, Journal of Cell Biology, 135(4), pp. 1097–1107. doi: 10.1083/jcb.135.4.1097.

Brugués, A. et al. (2014) ‘Forces driving epithelial wound healing’, Nature Physics, 10(9), pp. 683–690. doi: 10.1038/NPHYS3040.

Caglar, C. et al. (2021) ‘Fast transcriptional activation of developmental signalling pathways during wound healing of the calcareous sponge *Sycon ciliatum*’, bioRxiv, p. 2021.07.22.453456. doi: 10.1101/2021.07.22.453456

Carlson, B. (2007) Principles of Regenerative Biology. Elsevier.

Chen, T. et al. (2019) ‘Large-scale curvature sensing by directional actin flow drives cellular migration mode switching’, Nature Physics, 15(4), pp. 393–402. doi: 10.1038/s41567-018-0383-6.

Colgren, J. and Nichols, S. A. (2022) ‘MRTF specifies a muscle-like contractile module in Porifera’, Nature Communications, 13(1), pp. 1–11. doi: 10.1038/s41467-022-31756-9.

Cordeiro, J. V. and Jacinto, A. (2013) ‘The role of transcription-independent damage signals in the initiation of epithelial wound healing’, Nature Reviews Molecular Cell Biology. 14(4), pp. 249–262. doi: 10.1038/nrm3541.

Danjo, Y. and Gipson, I. K. (1998) ‘Actin “purse string” filaments are anchored by E-cadherin-mediated adherens junctions at the leading edge of the epithelial wound, providing coordinated cell movement’, Journal of Cell Science, 111(22), pp. 3323–3332. doi: 10.1242/jcs.111.22.3323.

Eerkes-Medrano, D. I. and Leys, S. P. (2006) ‘Ultrastructure and embryonic development of a syconoid calcareous sponge’, Invertebrate Biology, 125(3), pp. 177–194. doi: 10.1111/j.1744-7410.2006.00051.x.

Elliott, G. R. D. and Leys, S. P. (2007) ‘Coordinated contractions effectively expel water from the aquiferous system of a freshwater sponge’, Journal of Experimental Biology, 210(21), pp. 3736–3748. doi: 10.1242/jeb.003392.

Ereskovsky, A. et al. (2017) ‘Regeneration in white sea sponge *Leucosolenia complicata* (Porifera, Calcarea)’, Invertebrate Zoology, 41(2), pp. 108–113. doi: 10.15298/invertzool.14.2.02.

Ereskovsky, A. et al. (2019) ‘Mass mortality event of White Sea sponges as the result of high temperature in summer 2018’, Polar Biology, 42(12), pp. 2313–2318. doi: 10.1007/S00300-019-02606-0.

Ereskovsky, A. et al. (2021) ‘Whole-body regeneration in sponges: Diversity, fine mechanisms, and future prospects’, Genes, 12(4). doi: 10.3390/GENES12040506.

Ereskovsky, A. and Lavrov, A. (2021) ‘Porifera’, in Invertebrate Histology. Wiley, pp. 19–54. doi: 10.1002/9781119507697.ch2.

Ereskovsky, A. V. (2010) The comparative embryology of sponges, Springer Netherlands. doi: 10.1007/978-90-481-8575-7.

Ereskovsky, A. V. et al. (2015) ‘Oscarella lobularis (Homoscleromorpha, Porifera) Regeneration: Epithelial morphogenesis and metaplasia’, PLoS ONE, 10(8), pp. 1–28. doi: 10.1371/journal.pone.0134566.

Ereskovsky, A. V. et al. (2020) ‘Transdifferentiation and mesenchymal-to-epithelial transition during regeneration in Demospongiae (Porifera)’, Journal of Experimental Zoology Part B: Molecular and Developmental Evolution, 334(1), pp. 37–58. doi: 10.1002/jez.b.22919.

Fahey, B. and Degnan, B. M. (2010) ‘Origin of animal epithelia: Insights from the sponge genome’, Evolution and Development, 12(6), pp. 601–617. doi: 10.1111/j.1525-142X.2010.00445.x.

Farooqui, R. and Fenteany, G. (2005) ‘Multiple rows of cells behind an epithelial wound edge extend cryptic lamellipodia to collectively drive cell-sheet movement’, Journal of Cell Science, 118(1), pp. 51–63. doi: 10.1242/jcs.01577.

Fenteany, G., Janmey, P. A. and Stossel, T. P. (2000) ‘Signaling pathways and cell mechanics involved in wound closure by epithelial cell sheets’, Current Biology, 10(14), pp. 831–838. doi: 10.1016/S0960-9822(00)00579-0.

Gaino, E. and Magnino, G. (1999) ‘Dissociated cells of the calcareous sponge *Clathrina*: A model for investigating cell adhesion and cell motility in vitro’, Microscopy Research and Technique, 44(4), pp. 279–292.

Garcia-Fernandez, B. et al. (2009) ‘Epithelial resealing’, International Journal of Developmental Biology, 53(8–10), pp. 1549–1556. doi: 10.1387/ijdb.072308bg.

Gauthier M. (2009). Developing a sense of self: exploring the evolution of immune and allorecognition mechanisms in metazoans using the demosponge amphimedon queenslandica. PhD Thesis, School of Biological Sciences, The University of Queensland.

Gemberling, M. et al. (2013) ‘The zebrafish as a model for complex tissue regeneration’, Trends in genetics, 29(11), pp. 611–620. doi: 10.1016/J.TIG.2013.07.003.

Grasso, S., Hernández, J. A. and Chifflet, S. (2007) ‘Roles of wound geometry, wound size, and extracellular matrix in the healing response of bovine corneal endothelial cells in culture’, American Journal of Physiology - Cell Physiology, 293(4). doi: 10.1152/ajpcell.00001.2007.

Green, K. J. et al. (2020) ‘Tracing the Evolutionary Origin of Desmosomes’, Current Biology, 30(10), pp. R535–R543. doi: 10.1016/j.cub.2020.03.047.

Hay, E. D. and Zuk, A. (1995) ‘Transformations between epithelium and mesenchyme: normal, pathological, and experimentally induced’, American journal of kidney diseases: the official journal of the National Kidney Foundation, 26(4), pp. 678–690. doi: 10.1016/0272-6386(95)90610-X.

Jacinto, A. et al. (2000) ‘Dynamic actin-based epithelial adhesion and cell matching during *Drosophila* dorsal closure’, Current Biology, 10(22), pp. 1420–1426. doi: 10.1016/S0960-9822(00)00796-X.

Jankovics, F. and Brunner, D. (2006) ‘Transiently Reorganized Microtubules Are Essential for Zippering during Dorsal Closure in Drosophila melanogaster’, Developmental Cell, 11(3), pp. 375–385. doi: 10.1016/j.devcel.2006.07.014.

Jones, W. C. (1957) ‘The Contractility and Healing Behaviour of Pieces of Leucosolenia complicata’, Journal of Cell Science, s3-98(42), pp. 203–217. doi: 10.1242/jcs.s3-98.42.203.

Jones, W. C. (1966) ‘The structure of the porocytes in the calcareous sponge *Leucosolenia complicata* (Montagu)’, Journal of the Royal Microscopical Society, 85(1), pp. 53–62. doi: 10.1111/j.1365-2818.1966.tb02166.x.

Jonusaite, S., Donini, A. and Kelly, S. P. (2016) ‘Occluding junctions of invertebrate epithelia’, Journal of Comparative Physiology B: Biochemical, Systemic, and Environmental Physiology, 186(1), pp. 17–43. doi: 10.1007/s00360-015-0937-1.

Jopling, C., Boue, S. and Belmonte, J. C. I. (2011) ‘Dedifferentiation, transdifferentiation and reprogramming: Three routes to regeneration’, Nature Reviews Molecular Cell Biology, 12(2), pp. 79–89. doi: 10.1038/nrm3043.

Kamran, Z. et al. (2017) ‘In vivo imaging of epithelial wound healing in the cnidarian *Clytia hemisphaerica* demonstrates early evolution of purse string and cell crawling closure mechanisms’, BMC Developmental Biology, 17(1), pp. 1–15. doi: 10.1186/s12861-017-0160-2.

Kawamura, K. et al. (2008) ‘Multipotent epithelial cells in the process of regeneration and asexual reproduction in colonial tunicates’, Development, Growth & Differentiation, 50(1), pp. 1–11. doi: 10.1111/J.1440-169X.2007.00972.X.

Klarlund, J. K. (2012) ‘Dual modes of motility at the leading edge of migrating epithelial cell sheets’, Proceedings of the National Academy of Sciences of the United States of America, 109(39), pp. 15799–15804. doi: 10.1073/pnas.1210992109.

Korotkova, G. P. (1961). Regeneration and cellular proliferation in calcareous sponge *Leucosolenia complicata* Mont. Vestnik Leningrad University. 4 (21): 39–50.

Korotkova, G. P. (1963). Regeneration and somatic embryogenesis in calcareous sponges of the type Sycon. Vestnik Leningrad University. 3: 34–47.

Kretschmer, S. et al. (2017) ‘Imaging of Wound Closure of Small Epithelial Lesions in the Mouse Trachea’, American Journal of Pathology, 187(11), pp. 2451–2460. doi: 10.1016/j.ajpath.2017.07.006.

Kuipers, D. et al. (2014) ‘Epithelial repair is a two-stage process driven first by dying cells and then by their neighbours’, Journal of Cell Science, 127(6), pp. 1229–1241. doi: 10.1242/jcs.138289.

Kukulies, J., Naib-Majani, W. and Komnick, H. (1984) ‘Coincident filament distribution and histochemical localization of F-actin in the enterocytes of the larval dragonfly *Aeshna cyanea*’, Protoplasma, 121(1–2), pp. 157–162. doi: 10.1007/BF01279763.

Lane, N. J. and Flores, V. (1988) ‘Actin filaments are associated with the septate junctions of invertebrates’, Tissue and Cell, 20(2), pp. 211–217. doi: 10.1016/0040-8166(88)90042-0.

Lanosa, X. A. and Colombo, J. A. (2008) ‘Cell contact-inhibition signaling as part of wound-healing processes in brain’, Neuron Glia Biology, 4(1), pp. 27–34. doi: 10.1017/S1740925X09000039.

Lavrov, A. I. et al. (2018) ‘Sewing up the wounds: The epithelial morphogenesis as a central mechanism of calcaronean sponge regeneration’, Journal of Experimental Zoology Part B: Molecular and Developmental Evolution, 330(6–7), pp. 351–371. doi: 10.1002/jez.b.22830.

Lavrov, A. I. et al. (2020) ‘Intraspecific variability of cell reaggregation during reproduction cycle in sponges’, Zoology, 140. doi: 10.1016/J.ZOOL.2020.125795.

Lavrov, A. I. et al. (2022) ‘Fine details of the choanocyte filter apparatus in asconoid calcareous sponges (Porifera: Calcarea) revealed by ruthenium red fixation’, Zoology, 150, p. 125984. doi: 10.1016/j.zool.2021.125984.

Lavrov, A. I. and Kosevich, I. A. (2014) ‘Sponge cell reaggregation: Mechanisms and dynamics of the process’, Russian Journal of Developmental Biology, 45(4), pp. 205–223. doi: 10.1134/S1062360414040067.

Ledger, P. W. (1975) ‘Septate junctions in the calcareous sponge Sycon ciliatum’, Tissue and Cell, 7(1), pp. 13–18. doi: 10.1016/S0040-8166(75)80004-8.

Lee, J. S. Y. and Gotlieb, A. I. (2005) ‘Microtubules regulate aortic endothelial cell actin microfilament reorganization in intact and repairing monolayers’, Histology and Histopathology, 20(2), pp. 455–465.

Leys, S. P. and Hill, A. (2012) The Physiology and Molecular Biology of Sponge Tissues. 1st edn, Advances in Marine Biology. doi: 10.1016/B978-0-12-394283-8.00001-1.

Leys, S. P., Nichols, S. A. and Adams, E. D. M. (2009) ‘Epithelia and integration in sponges’, Integrative and Comparative Biology, 49(2), pp. 167–177. doi: 10.1093/icb/icp038.

Leys, S. P. and Riesgo, A. (2012) ‘Epithelia, an evolutionary novelty of metazoans’, Journal of Experimental Zoology Part B: Molecular and Developmental Evolution, 318(6), pp. 438–447. doi: 10.1002/jez.b.21442.

Martin, P. and Lewis, J. (1992) ‘Actin cables and epidermal movement in embryonic wound healing’, Nature, 360(6400), pp. 179–183. doi: 10.1038/360179a0.

Miller, P. W. et al. (2018) ‘Analysis of a vinculin homolog in a sponge (phylum Porifera) reveals that vertebrate-like cell adhesions emerged early in animal evolution’, Journal of Biological Chemistry, 293(30), pp. 11674–11686. doi: 10.1074/jbc.RA117.001325.

Mitchell, J. M. and Nichols, S. A. (2019) ‘Diverse cell junctions with unique molecular composition in tissues of a sponge (Porifera)’, EvoDevo, 10(1), pp. 1–16. doi: 10.1186/s13227-019-0139-0.

Nickel, M. et al. (2011) ‘The contractile sponge epithelium sensu lato-body contraction of the demosponge *Tethya wilhelma* is mediated by the pinacoderm’, Journal of Experimental Biology, 214(10), pp. 1692–1698. doi: 10.1242/jeb.049148.

Nier, V. et al. (2015) ‘Tissue fusion over nonadhering surfaces’, Proceedings of the National Academy of Sciences of the United States of America, 112(31), pp. 9546–9551. doi: 10.1073/pnas.1501278112.

Nobes, C. D. and Hall, A. (1995) ‘Rho, Rac, and Cdc42 GTPases regulate the assembly of multimolecular focal complexes associated with actin stress fibers, lamellipodia, and filopodia’, Cell, 81(1), pp. 53–62. doi: 10.1016/0092-8674(95)90370-4.

Nusrat, A., Delp, C. and Madara, J. L. (1992) ‘Intestinal epithelial restitution characterization of a cell culture model and mapping of cytoskeletal elements in migrating cells’, Journal of Clinical Investigation, 89(5), pp. 1501–1511. doi: 10.1172/JCI115741.

Padua, A. and Klautau, M. (2016) ‘Regeneration in calcareous sponges (Porifera)’, Journal of the Marine Biological Association of the United Kingdom, 96(2), pp. 553–558. doi: 10.1017/S0025315414002136.

Pang, S. C., Daniels, W. H. and Buck, R. C. (1978) ‘Epidermal migration during the healing of suction blisters in rat skin: A scanning and transmission electron microscopic study’, American Journal of Anatomy, 153(2), pp. 177–191. doi: 10.1002/aja.1001530202.

Pavans de Ceccatty, M. (1981) ‘Demonstration of actin filaments in sponge cells’, Cell Biology International Reports, 5(10), pp. 945–952. doi: 10.1016/0309-1651(81)90210-1.

Pavans de Ceccatty, M. (1986) ‘Cytoskeletal organization and tissue patterns of epithelia in the sponge Ephydatia mülleri’, Journal of Morphology, 189(1), pp. 45–65. doi: 10.1002/jmor.1051890105.

Pedersen, K. J. (1964) ‘The cellular organization of *Convoluta convoluta*, an acoel turbellarian: A cytological, histochemical and fine structural study’, Zeitschrift für Zellforschung und Mikroskopische Anatomie, 64(5), pp. 655–687. doi: 10.1007/BF01258542.

Peña, J. F. et al. (2016) ‘Conserved expression of vertebrate microvillar gene homologs in choanocytes of freshwater sponges’, EvoDevo, 7(1), pp. 1–15. doi: 10.1186/s13227-016-0050-x.

Pope, K. L. and Harris, T. J. C. (2008) ‘Control of cell flattening and junctional remodelling during squamous epithelial morphogenesis in Drosophila’, Development, 135(13), pp. 2227–2238. doi: 10.1242/dev.019802.

Reffay, M. et al. (2014) ‘Interplay of RhoA and mechanical forces in collective cell migration driven by leader cells’, Nature Cell Biology, 16(3), pp. 217–223. doi: 10.1038/ncb2917.

Richardson, R. et al. (2016) ‘Re-epithelialization of cutaneous wounds in adult zebrafish combines mechanisms of wound closure in embryonic and adult mammals’, Development, 143(12), pp. 2077–2088. doi: 10.1242/dev.130492.

Simion, P. et al. (2017) ‘A Large and Consistent Phylogenomic Dataset Supports Sponges as the Sister Group to All Other Animals’, Current Biology, 27(7), pp. 958–967. doi: 10.1016/j.cub.2017.02.031.

Simpson, T. L. (1984) Cell Biology of Sponges, Springer.

Smith, C. L. et al. (2014) ‘Novel cell types, neurosecretory cells, and body plan of the early-diverging metazoan Trichoplax adhaerens’, Current Biology, 24(14), pp. 1565–1572. doi: 10.1016/j.cub.2014.05.046.

Smith, C. L. and Reese, T. S. (2016) ‘Adherens junctions modulate diffusion between epithelial cells in Trichoplax adhaerens’, Biological Bulletin, 231(3), pp. 216–224. doi: 10.1086/691069.

Stramer, B. et al. (2005) ‘Live imaging of wound inflammation in *Drosophila* embryos reveals key roles for small GTPases during in vivo cell migration’, Journal of Cell Biology, 168(4), pp. 567–573. doi: 10.1083/jcb.200405120.

Tamada, M. et al. (2007) ‘Two distinct modes of myosin assembly and dynamics during epithelial wound closure’, Journal of Cell Biology, 176(1), pp. 27–33. doi: 10.1083/jcb.200609116.

Tang, W. J. et al. (2022) ‘Single-cell resolution of MET-and EMT-like programs in osteoblasts during zebrafish fin regeneration’, iScience, 25(2), p. 103784. doi: 10.1016/J.ISCI.2022.103784.

Tétreault, M. P. et al. (2005) ‘Differential growth factor induction and modulation of human gastric epithelial regeneration’, Experimental Cell Research, 306(1), pp. 285–297. doi: 10.1016/j.yexcr.2005.02.019.

Thiemann, M. et al. (1989) ‘Microfilaments and microtubules in isolated fiber cells of *Trichoplax adhaerens* (Placozoa)’, Zoomorphology, 109, pp. 89–96.

Tiozzo, S. and Copley, R. R. (2015) ‘Reconsidering regeneration in metazoans: An evo-devo approach’, Frontiers in Ecology and Evolution, 3, pp. 1–12. doi: 10.3389/fevo.2015.00067.

Tyler, S. and Hooge, M. (2004) ‘Comparative morphology of the body wall in flatworms (Platyhelminthes)’, Canadian Journal of Zoology, 82(2), pp. 194–210. doi: 10.1139/z03-222.

Valisano, L. et al. (2006) ‘Primmorphs formation dynamics: a screening among Mediterranean sponges’, Marine Biology, 149(5), pp. 1037–1046. doi: 10.1007/s00227-006-0297-1.

Vedula, S. R. K. et al. (2014) ‘Epithelial bridges maintain tissue integrity during collective cell migration’, Nature Materials, 13(1), pp. 87–96. doi: 10.1038/nmat3814.

Vedula, S. R. K. et al. (2015) ‘Mechanics of epithelial closure over non-adherent environments’, Nature Communications, 6, pp. 1–10. doi: 10.1038/ncomms7111.

Vervoort, M. (2011) ‘Regeneration and Development in Animals’, Biological Theory, 6(1), pp. 25–35. doi: 10.1007/S13752-011-0005-3.

Vijayakumar, S. et al. (1999) ‘Hensin remodels the apical cytoskeleton and induces columnarization of intercalated epithelial cells: Processes that resemble terminal differentiation’, Journal of Cell Biology, 144(5), pp. 1057–1067. doi: 10.1083/jcb.144.5.1057.

Wachtmann, D., Stockem, W. and Weissenfels, N. (1990) ‘Cytoskeletal organization and cell organelle transport in basal epithelial cells of the freshwater sponge *Spongilla lacustris*’, Cell and Tissue Research, 261(1), pp. 145–154. doi: 10.1007/BF00329447.

Waterman-Storer, C. M., Salmon, W. C. and Salmon, E. D. (2000) ‘Feedback interactions between cell-cell adherens junctions and cytoskeletal dynamics in newt lung epithelial cells’, Molecular Biology of the Cell, 11(7), pp. 2471–2483. doi: 10.1091/mbc.11.7.2471.

Yin, J., Lu, J. and Yu, F.-S. X. (2008) ‘Role of small GTPase Rho in regulating corneal epithelial wound healing’, Investigative Opthalmology & Visual Science, 49(3), p. 900–909. doi: 10.1167/iovs.07-1122.

Zahm, J. M. et al. (1997) ‘Cell migration and proliferation during the in vitro wound repair of the respiratory epithelium’, Cell Motility and the Cytoskeleton, 37(1), pp. 33–43.

Zahm, J. M., Chevillard, M. and Puchelle, E. (1991) ‘Wound repair of human surface respiratory epithelium.’, American journal of respiratory cell and molecular biology, 5(3), pp. 242–248. doi: 10.1165/ajrcmb/5.3.242.

