## Supplementary figures and images for "Regeneration in calcareous sponge relies on ‘purse-string’ mechanism and the rearrangements of actin cytoskeleton"

### Figure S1

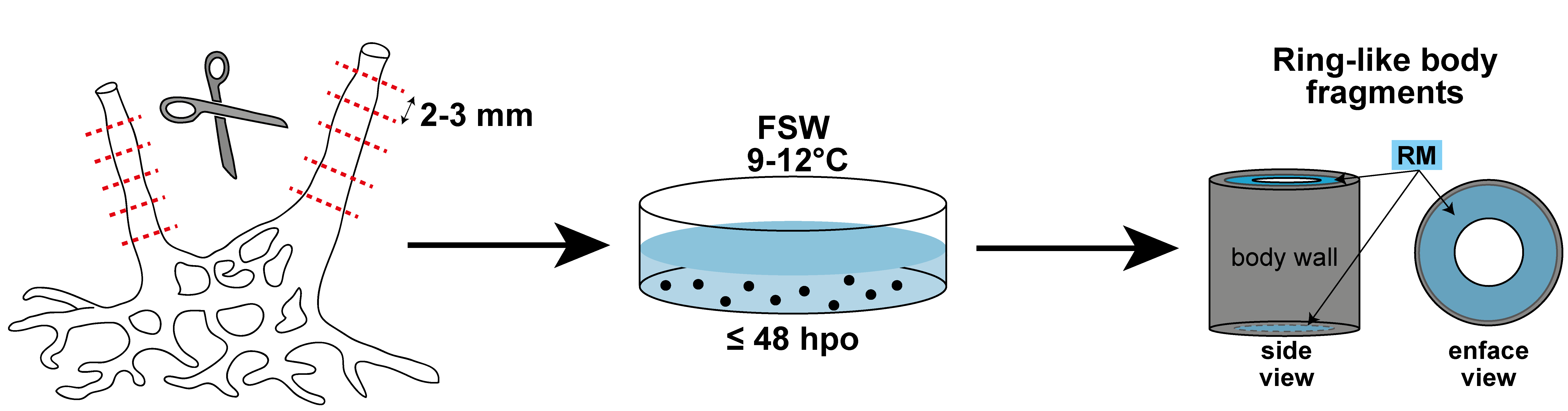
